# Predicting cell type-specific epigenomic profiles accounting for distal genetic effects

**DOI:** 10.1101/2024.02.15.580484

**Authors:** Alan E Murphy, William Beardall, Marek Rei, Mike Phuycharoen, Nathan G Skene

**Affiliations:** UK Dementia Research Institute at Imperial College London, London W12 0BZ, UK; Department of Brain Sciences, Imperial College London, London W12 0BZ, UK; Department of Bioengineering, Imperial College London, London SW7 2AZ, UK; Department of Computing, Imperial College London, London SW7 2RH, UK; Division of Informatics, Imaging & Data Sciences, University of Manchester, M13 9PL, UK

**Author notes:** Correspondence should be addressed to A.M. (email: Alan Murphy) and N.S. (email: Nathan Skene).

## Abstract

Understanding genetic variants’ effects on the epigenome is crucial for interpreting genome-wide association studies (GWAS) results, yet profiling these effects across the non-coding genome remains challenging due to the scalability limits of experimental methods. This necessitates accurate computational models. Existing machine learning approaches, while progressively improving, are confined to the cell types they were trained on, limiting their applicability. Here, we propose the development of a deep learning model, Enformer Celltyping, which can both incorporate distal effects of DNA interactions, up to 100,000 base-pairs away, and predict epigenetic signals in a cell type-agnostic fashion. We demonstrate that Enformer Celltyping’s predictions of the epigenome out-perform the current best-in-class approach, have strong performance in a range of different cell types and biological regions and generalise to cell types assayed independently of the original training set. Finally, we propose an approach to test epigenetic models’ performance on genetic variant effect predictions using regulatory quantitative trait loci mapping studies, highlighting Enformer Celltyping’s and other genomic deep learning models’ limitations for this task. Our work introduces a new, accurate model for cell type-agnostic histone mark predictions and highlights the benefit of transfer learning for more efficient model development for deep learning in genomics. We make our customisable and efficient transfer learning approach for Enformer available to the community.

## Background

Recent large-scale genetic efforts, genome-wide association studies (GWAS), have identified associated variants for a myriad of diseases^1,2,3^. However, these often lie in non-coding, regulatory regions and cannot be associated with any functional outcomes. For example, recent GWAS have found more than 90% of Single Nucleotide Polymorphisms (SNPs) reside in non-coding regions^4^. One of the main challenges to understanding the function of these regulatory variants is that gene regulatory mechanisms are highly cell type-specific^5^. Genetic sequence variants which are associated with the function or activity of regulatory elements, quantitative trait loci (QTLs), will tend to exert their effects in a cell type-specific manner. Thus, understanding the cell type-specific effect of genetic variants in epigenetic regulation would help identify the biological processes they act on.

Mapping out molecular and regulatory QTLs comprehensively in disease-relevant cell types would enable the interpretation of functional outcomes of genetic variants on gene expression and regulation. Techniques to study QTLs include CRISPR interference (CRISPRi)^6^ which is used to perturb regulatory regions and measure the corresponding impact and Massively Parallel Reporter Assays (MPRAs)^7^ to interrogate multiple candidate genetic regulatory elements and their activity. However, these approaches lack the *in vivo* capabilities or scalability to large sample sizes thus limiting their efficacy. Currently, the most suitable approach is to conduct population studies measuring individuals’ genomic variance and its correlation to regulatory elements, i.e. xQTL mapping studies. QTL mapping of regulatory elements is distinguishable to that of gene expression (eQTL mapping studies) which has gained a lot more attention, including full catalogues of studies with sample sizes in the high hundreds^8^ and the largest eQTL to date having more than 30,000 samples^9^. Some QTL regulatory mapping studies of peripheral tissues and cell types, where samples can be collected much less invasively, have been conducted. Notably, Kundu and colleagues^10^ comprehensively profiled genetic and epigenetic variation in three major human immune cell types (CD14+ monocytes, CD16+ neutrophils and naive CD4+T cells). Using these data, they report the largest QTL study of blood cell types to date and linked regulatory QTLs to non-coding genetic risk variants for immune disorders^10^. However, despite being the largest epigenetic QTL to date, this study had a sample size of just 196, far less than the average expression QTL study. Moreover, the study is one of just a small number of histone mark (hQTL) or transcription factor (tfQTL) studies conducted. Robustly identifying regulatory QTLs from primary data in QTL regulatory mapping studies requires very large sample numbers. This is due to the high-dimensional nature of the association involving pairwise testing for associations between all SNPs and all regulatory elements. Undertaking such studies in large sample sizes and on every possible cell type of interest is impractical.

An alternative is to predict the effect of genetic variants using machine learning approaches, known as *in silico* mutagenesis. Numerous machine learning approaches have been developed to predict the effect of genetic variants across different tasks such as transcription factor binding^11,12^, epigenetic marks^13^, gene expression^14^ profiles, exon splicing locations^15^ or even topologically associating domains (TADs) boundaries^16^. One of the first of these approaches was DeepSea^17^. DeepSea employed a deep learning framework a convolutional neural network (CNN), to learn transcription factor binding profiles directly from genetic sequence. The model was trained on wild type profiles and the effect of genetic variants were predicted after model training, by comparing the wild type prediction against that based on the DNA sequence with the imputed SNP. While most subsequent models have followed this approach, one notable exception is Wang *et al.*^18^, who trained their model directly on the genetic variant information. The authors first identified the causal SNP from those that were in linkage disequilibrium (LD) through a statistical fine-mapping approach; FINEMAP^19^, to avoid the model learning spurious effects of genetic variants. Although this novel approach showed promise, recent reviews have highlighted that despite considerable advancements, there is still work to be done to improve current fine-mapping approaches^20,21,22^.

The main advancement of more recent approaches in the field has focused on the amount of DNA sequence which models can take into account simultaneously to make the prediction. This has generally led to greater performance of these models on held-out genomic positions. Intuitively, these larger window sizes mean that when predicting the profile at a given location, distal effects of DNA sequence can be modelled. The maximum distance from the predicted profile to DNA sequence in either direction of such models has increased from 500 base-pairs for DeepSea^17^ to 20,000 base-pairs in Basenji2^13^ and in the current best-in-class approach, Enformer^23^, to 100,000 base-pairs. Enformer swapped out convolutional layers from Basenji2’s original architecture with multi-headed attention layers and three custom, relative positional encoding functions to dramatically increase the receptive field.

Importantly, all three of the aforementioned models were trained solely on DNA sequence and thus cannot predict cell type profiles that were not seen in the training phase. Their use is limited to the prediction of the effect of genetic variants in cell types included in training. Although valuable, it would be beneficial to predict epigenetic profiles and the effect on these from genetic variants in previously unseen cell types where collecting cell type epigenetic profiles can be invasive, time-consuming and costly. Some past models have attempted to generalise predictions of epigenetic profiles to other cell types by using cell type-specific chromatin accessibility information, often Dnase-Seq or ATAC-Seq, such as Leopard^12^, Catchitt^11^ (both predict transcription factor binding profiles) and Epitome^24^ (histone mark profiles). Notably, these approaches do not consider distal genetic information, instead using local genetic windows, far shorted than that of Enformer, and other local features to the infer the cell type variability. However, this local information may not be enough for a model to fully capture cell type representations. An alternative approach of cell type embedding has been explored in Avocado^25^. Avocado, a deep neural network tensor factorisation method, is designed to impute missing epigenetic marks and transcription factor binding profiles based on the profiles present for the cell type. Avocado achieves this by embedding substantial window sizes of these profiles (>1 million base-pairs) along with the specific cell type and assay, to make these imputations. This embedding is similar to the word embedding techniques used in Natural Language processing (NLP)^26^.

Although multiple approaches have now been developed for *in silico* mutagenesis, there is little consensus on how best to benchmark performance. Experimental datasets including CRISPR, MPRAs and xQTL mapping studies have all been proposed^23,27,28^. However, similar to the restrictions for using these approaches to experimentally measure the effect of genetic variants, currently only xQTL mapping studies can be carried out *in vivo* on large sample sizes. Thousands of genetic variants are tested in one study based on the natural variation in the population. However, associations detected in xQTL mapping studies are confounded by linkage disequilibrium (LD), where alleles are co-inherited based on their physical proximity^29^, making it difficult to identify the causal SNPs^30^. Therefore, LD needs to be taken into account when comparing these datasets against model predictions. Moreover, as previously mentioned, the largest regulatory QTL mapping studies have been conducted on histone marks thus we use the prediction of histone marks and the effect of genetic variants on these regulatory signals as the first step to understanding all classes of epigenetic regulation like transcription factor binding or DNA methylation.

Here, we propose Enformer Celltyping, a self-attention based neural network model to predict histone mark activity in previously unseen cell types. Histone mark profiles are highly cell type-specific^31^ and changes in their activity have been investigated in a multitude of cell processes including aging^32^, transcription factor activity^33^ and disease^34^. However, experimentally measuring effects of genetic variants on histone marks in cell types of interest is experimentally impractical. Thus, being able to predict these marks in a cell type of interest is paramount to elucidate the cell’s epigenetic regulation.

Our model predicts six histone mark profiles from DNA sequence and chromatin accessibility information and is trained on 104 ENCODE samples, hereafter referred to as cell types despite containing tissue samples, isolated cell types and cell lines, sourced from EpiMap^35^. We use a custom transfer learning approach on a pre-trained Enformer^23^ model to account for the distal effect of genetic code on these regulatory signals. This approach results in substantial memory and time savings during model training. Enformer Celltyping also embeds a genome-wide and local representation of cell types from chromatin accessibility information to extend Enformers’ capabilities, making it applicable to any cell type of interest where accessibility data is available. This embedding is similar to that of Avocado’s^25^, taking inspiration from the field of NLP. We validate Enformer Celltyping’s performance against best-in-class approaches and in a number of differing biological settings, inspect the cell type-specific information the model learns and apply it to predict the effect of genetic variants on histone mark profiles. Moreover, we outline an approach to validate the predictive performance of epigenetic models at genetic variant effect predictions using QTL studies with known associations and accounting for LD. We have made a pre-trained version of Enformer Celltyping available along with a fine-tuneable version of the Enformer model, something which was previously not available, so others may make use of these in their research (code: https://github.com/neurogenomics/EnformerCelltyping and pre-trained model: https://figshare.com/account/home#/projects/159143).

## Methods

### Data collection and processing

The training data was derived from GRCh37 DNA sequence and p-value continuous tracks after peak calling in EpiMap^35^, encompassing data from the ENCODE project^31^. The DNA sequence was one-hot encoded following the same approach as previous models^17,12,13,23^. The aim for the histone mark tracks data collection from EpiMap was to maximise both the number of histone marks the model could predict and also the number of cell types which due to the sparse nature of ENCODE data, results in a trade-off between the two (Additional file 1: Fig. S9). Any cell types with assays marked as low quality by EpiMap were removed from the training set. The final training set included ChIP-seq histone mark data for H3K27ac, H3K4me1, H3K4me3, H3K9me3, H3K27me3 and H3K36me3 from 104 cell types (Additional file 2: Table S1). Given the current preference of ATAC-seq over DNase-Seq to measure chromatin accessibility (ATAC-Seq has similar sensitivity and specificity to DNase-Seq with three to five times fewer cells^36^), we opted to train our model using ATAC-seq data so it would be more applicable to future users. To avoid losing cell types, imputed chromatin accessibility ATAC-Seq data sourced from EpiMap^35^ was used for any where there was not an observed track which was the case for the majority (98 out of 104) of cell types. All tracks were converted from 25 base-pair average signals to 128 base-pair averaging to match the resolution of the Enformer^23^. All training data tracks measure the −log_10_ adjusted p-value from MACS2 peak calling^37^, indicating the statistical significance of a genomic position i.e. is the site likely to be a real binding site when compared against a background. Despite EpiMap’s uniform reprocessing, to avoid possible issues like sequencing depth biases, we followed previous approaches of training on the arcsinh-transformed signal^25,38,39^:

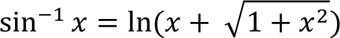

We achieved clear separation across epigenetic marks on our training cell types after this transformation which was not present otherwise (Additional file 1: Fig. S10). The average chromatin accessibility and histone mark signals, used for model training, were derived by averaging these 104 cell types’ tracks (see Methods section Enformer Celltyping architecture).

Two separate sets of cells were used as test cases for the model; 3 immune cell types sourced from EpiMap^35^ (CD14+ monocytes, CD16+ neutrophils and naive CD4+T cells) and Nott *et al.*’s^40^ isolated cell types from resected cortical brain tissue (PU.1+ microglia, NeuN+ neuronal, OLIG2+ oligodendrocyte and NeuN-LHX2+ astrocytes). The Nott *et al*., data was processed to produce consensus peak p-value tracks across samples (different individuals were taken as separate replicates) using the ENCODE ChIP-Seq and ATAC-Seq pipelines (https://github.com/ENCODE-DCC/chip-seq-pipeline2 and https://github.com/ENCODE-DCC/atac-seq-pipeline) with the supplied conda environment and default parameters. One exception to the default parameters was subsampling the signals to 30 million reads for the ChIP-Seq and 50 million for the ATAC-Seq to match the processing of the EpiMap data and avoiding performance loss due to sequencing depth issues^41^. Identically to the training cell types, the test sets were averaged to 128 base-pair resolution and arcsinh-transformed. However, none of these test cell types or any similar cell types, such as other immune cells, from EpiMap were used in the training set. Moreover, these were not used to derive the average histone mark or chromatin accessibility tracks to avoid data leakage between training and test sets^41^.

The histone QTL (hQTL) Blueprint phase 2 data^10^ was downloaded from the European Genome-Phenome Archive (ID: EGAD00001005199). The six hQTL sets (CD14+ monocytes, CD16+ neutrophils and naive CD4+T cells for histone marks H3K4me1 and H3K27ac) were uniformly processed using MungeSumstats (v1.6.0)^42^, using the format_sumstats function and default parameters with the exception of the check_dups parameter which was set to false to enable quality control and formatting of QTL datasets. To transform the hQTL datasets to be compared with Enformer Celltyping’s SNP predictions, all SNP to histone mark peak interactions further than Enformer Celltyping’s receptive field, or any trans interactions, were removed along with all SNPs not included in HapMap3^30^. SNPs effects were then aggregated to a single value per SNP using the following transformation, as done in previous work^23,43,30^:

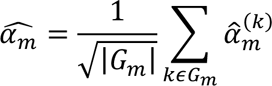

Where *G*_*m*_ is the set of all histone mark peaks for which a cis-hQTL test was performed for variant *m* and 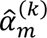 is the marginal correlation of SNP *m* and histone mark peak *k*’s signal^30^ (i.e. the Beta effect value from the study).

### Calculating Genetic Variant distance from epigenetic effect peak in QTL associations

The distance from the SNP to the associated histone mark peak for the six histone mark hQTL BLUEPRINT Phase 2 study^10^ were aggregated to one genetic variant per LD block to avoid double counting. Aggregation was done to keep SNPs with the lowest p-value for each LD block and if multiple with the same, the shortest distance to the effect region (to avoid biases towards distal interactions). The Pickrell LD blocks^44^ (available from https://bitbucket.org/nygcresearch/ldetect-data) were used for this analysis (Additional file 1: Fig. S2). This highlights the need for models which can account for these distal interactions when predicting epigenetic profiles, not only transcriptional profiles.

### Training Data

Enformer Celltyping was trained on 67,007 GRCh37 genomic regions across chromosomes 1-22 and across the 104 training cell types. The regions were picked by first binning the genome based on the size of the model’s predictive window (114,688 base-pairs). These bins were then filtered at a DNA level and also a cell type level. The DNA filters were to exclude bins in ENCODE blacklist regions^31^ and ensuring subsequent genomic positions up and downstream of the region so the model can utilise the full input window size (196,608 base-pairs). The cell type level filters were to ensure a coverage of histone mark signals > 12.5% in the region (based on a −log_10_ p-value cut-off of 2 to mark a peak^35^ for a mark in a 128 base-pair bin). This was done for each histone mark independently and then all histone mark regions were down-sampled to match that of the histone mark with the lowest number of peaks, resulting in 11,903 positions for each histone mark. This approach ensured equal representation and avoided the model prioritising training on a subset of over-represented marks^41^. The filtering resulted in 67,007 genomic regions across all training cell types and corresponds to 14,188 unique DNA positions which were split into a training and validation set by chromosome, a similar number of unique genomic regions to past approaches^13,23^. The forward and reverse complement sequence and small, random sequence shifts up and downstream for each were also generated for each genomic region to be included in the training set. The validation set was created by splitting 20% of the training cell types and the start position for these was randomly permuted around the chosen regions to avoid the model overfitting to the training bins. A list of the training positions and cell types is available for download through figshare (https://figshare.com/account/home#/projects/159143).

For each position in each histone mark, the average and distribution of arcsinh-transformed, −log_10_ p-values across the training cell types were calculated for the pre-training step (see Methods section Enformer Celltyping architecture). This distribution was created by binning signals for each histone mark and genomic region, based on their −log_10_ p-value signal into 10 groups at 0.5 intervals from 0 to 4.5. The distributions for each histone mark were calculated before model training, the code for which (and all other data processing and model training/testing) is available in our Github repository (https://github.com/neurogenomics/EnformerCelltyping).

### Pre-trained Enformer model

Enformer Celltyping uses a pre-trained version of Enformer^23^ which has had its pre-trained layers removed after the multi-headed attention layers and its weights frozen (Additional file 1: Fig. S1). A custom approach was developed to be able to apply transfer learning to Enformer. The Enformer model available from Tensorflow Hub has not been formatted to allow modifying of its layers or unfreezing weights (it doesn’t expose a *call* method, or any of its internal variables, and lacks a model signature). To circumvent this issue, we first extracted the weight matrices from the available, trained version of Enformer and rebuilt an untrained version of Enformer following the architecture provided in the original study^23^. We then manually mapped layers across the two models and once completed, updated the weights in each layer of the untrained Enformer model to the values of the trained version of Enformer. Once complete, we tested the predictions against the trained version of Enformer to ensure the weights were mapped correctly. From here, we removed the final layers after the multi-headed attention layers and froze the weights of the other layers and incorporated this pre-trained, chopped version of Enformer into our Enformer Celltyping model. This recreated version of Enformer is now fully customisable allowing flexibility in the transfer learning approach such as fine-tuning through Tensorflow API. We have made an automated script to create this version of Enformer available in our Github repository (https://github.com/neurogenomics/EnformerCelltyping) so others can similarly use Enformer in their own problem domains.

### Enformer Celltyping architecture

Enformer Celltyping was implemented in Tensorflow v2.4 with Sonnet v2 functionality in the pre-trained Enformer model. Enformer Celltyping was trained with a two-step approach – a pre-training step with a separate training of sequence and accessibility submodules, and a subsequent cell type-specific training step, combining the output of the two model branches. The pre-training step improved model performance overall (Additional file 1: Fig. S4).

For the pre-training, Enformer Celltyping’s architecture is split into two separate submodules (Additional file 1: Fig. S1a), both with their own optimiser and loss function. The first “DNA” submodule uses DNA sequence to predict the average and distribution of each histone mark. The second “celltyping” submodule predicts the difference between the average histone mark signal and the cell type-specific signal. These are explained in more detail below; how the data is constructed is discussed in Methods section Training data. The output blocks of the first DNA submodule is shown in Additional file 1: Fig S1a as “Output avg” and “Output distribution” while the second celltyping submodule’s pretraining output is shown as “Output delta”. It is the first “DNA” submodule which uses the pre-trained, chopped, frozen version of Enformer^23^ through a transfer learning approach (discussed above in “Pre-trained Enformer model”).

The first is a “DNA” submodule takes one-hot encoded DNA sequence of length 196,608 base-pairs (A = [1,0,0,0], C = [0,1,0,0], G = [0,0,1,0], T = [0,0,0,1]) as its input, and processes it through four main parts: (1) seven convolutional blocks with pooling, (2) eleven transformer blocks, and (3) a cropping layer and (4) a further dense layer and convolutional block and output layer. During both training steps, the layers corresponding to the first three parts of the architecture (the pre-trained Enformer layers) were frozen and only the final part’s weights were updated through backpropagation. The DNA submodule outputs an average cell type prediction and distribution across the training cell types for each histone mark at the genomic region. The average cell type −log10 p-value score for each histone mark, used to train the model, was derived from the 104 EpiMap, training cell types (see Methods section Data collection and processing). The distribution data corresponds to the proportion of training cells whose histone mark, arcsinh-transformed −log_10_ p-value fall into each of 10 bins between 0 and >=4.5 at 0.5 intervals for a given genomic location. For example, if for H3K27ac signal at a genomic location, all cell types have no signal i.e. −log_10_ p-value = 0, then the distribution to predict would be [**1**,0,0,0,0,0,0,0,0,0,0]. Whereas, if two cell types of the 104 had −log_10_ p-value = 5.5, then the distribution would be [**.98**,0,0,0,0,0,0,0,0,0,**0.019**]. Including the distribution of histone mark signals as well as the average in the DNA module prediction for the pre-training step helps capture variability of regions across cell types and improved the overall performance of Enformer Celltyping (Additional file 1: Fig. S5).

The second “celltyping” submodule takes two inputs, a local and global representation of chromatin accessibility (ATAC-Seq) for the cell type. The local chromatin accessibility is the ATAC-Seq from the 199,936 base-pairs averaged at 128 base-pair resolution of the same genomic location as the DNA (1,562 positions). This local chromatin accessibility signal is first pre-processed by calculating the difference between the cell type-specific signal and the average chromatin accessibility signal of the training before passing to the model. Whereas, the global chromatin accessibility corresponds to the chromatin accessibility for 3,000 base-pairs around the transcriptional start site of 1,216 marker genes, averaged at 250 base-pair resolution (3.648 million base-pairs, input size of 14,592 positions). These marker genes were taken from a database of all known human marker genes derived from over 1,000 single-cell RNA-Seq experiments for any cell type, collated by PangloaDB^45^. Human marker genes were excluded if they fell outside of chromosomes 1-22 or in ENCODE blacklist regions^31^, leaving 1,216 genes. The list of PangloaDB genes and code to derive the global chromatin accessibility is available at: https://github.com/neurogenomics/EnformerCelltyping. We tested the performance of using each of the global and local chromatin accessibility signals separately and combined for a subset of 300 training steps, highlighting the benefit of incorporating both (Additional file: Fig. S3). Note that the global chromatin accessibility signal will be the same for a cell type regardless of the genomic region it is predicting in. Similar to Avocado’s approach, the global and local chromatin accessibility information were embedded at different resolutions to capture differing information about the epigenetic landscape^25^. These embeddings are concatenated and passed through two dense layers before being passed to a separate output head per histone mark. The celltyping submodule outputs the difference between the average histone mark prediction and the cell type of interest for the genomic region.

For the full training step, Enformer Celltyping combines the three outputs; “Output avg”, “Output distribution” and “Output delta”, of the “DNA” and “celltyping” submodules with subsequent convolutional and dense layers (Additional file 1: Fig. S1b) to predict the cell type-specific histone mark signal for six histone mark (H3K27ac, H3K4me1, H3K4me3, H3K9me3, H3K27me3 and H3K36me3), in a given genomic region. Enformer Celltyping’s output has six channels, one for each histone mark and 896 positions, corresponding to the centre 114,688 base-pairs of the input window, aggregated into 128 base-pair resolution bins. Only the centre 114,688 base-pairs are predicted on to avoid predicting on the edge positions which do not have as many neighbouring positions upon which to make a prediction^23^. Enformer Celltyping is trained to predict arcsinh-transformed −log_10_ p-value, however, this can be converted back to a raw −log_10_ p-value which Enformer Celltyping outputs by default. All subsequent analysis was done using the raw −log_10_ p-value signal unless otherwise stated.

### Enformer Celltyping Training

Enformer Celltyping was trained using Tensorflow API for 1,000 steps of a batch size of 128 cell type-specific genomic positions (128,000 positions) as a pre-training stage taking approximately 4 days (see Methods section Enformer Celltyping architecture). The combined architecture, full training stage was run for 4 full epochs (6,940 steps with a batch size of 128 cell type-specific genomic positions equating to 888,320 positions) stopping when the model started to overfit on the validation dataset, taking approximately 1.5 days on a Nvidia A100, 80GB RAM GPU. A learning rate of 0.0002 was used for pre-training and 0.005 for the full training stage and the ADAM optimiser^46^ was used for both. The low initial learning rate was chosen to allow for learning rate warmup and avoid early overfitting^47^ after which we matched the learning rate to similar models^23,14^. In the pre-training stage, the loss function differed across the submodules; a Poisson negative log-likelihood loss function was used for the average signal prediction (the same as past approaches^13,23^) and cross-entropy loss was used for the distribution in the DNA submodule whereas a mean squared error (MSE) loss function was used for the celltyping submodule following other epigenetic embedding approaches^25,5^, given the possibility of negative values. Since Enformer Celltyping freezes the weights relating to the chopped, pre-trained Enformer model (see Methods section Pre-trained Enformer model), the output from these layers for each genomic position was computed just once for each genomic location and then load as needed for each epoch. As a result of this, training was substantially quicker than for Enformer in the original study^23^ despite our model seeing more positions for multiple epochs compared to Enformer’s single epoch approach, totalling 132 GPU hours for pre and full training stages versus 5,376 GPU hours.

### Testing receptive field

To test the receptive field of both Enformer and Enformer Celltyping, we evaluated the effect of random genetic variants on histone mark predictions across increasing distances (from 0 to 98,304 base-pairs up and downstream). Specifically, we measured the average change in prediction across reference and alternative sequence on the centre four positions (512 base-pairs) of the output signal. DNA sequences were simulated by randomly sampling base-pairs. Next, a single random genetic variant per simulated DNA sequence was inserted and the effect on prediction measured. The location of the genetic variants were spread across 1,000 evenly spaced locations across the 196,608 base-pair input. This random sequence and random variant simulation was repeated 100 times for each genetic variant location (i.e. 100,000 simulations overall) to capture the average effect of any variant. All six output channels of Enformer Celltyping were averaged whereas only the output channels from Enformer corresponding to histone mark signals were considered. The same random iteration seed was used for both models so that the same DNA sequences and genetic variants would be simulated. The average change across reference and alternative was shown at the position of the genetic variant (Fig. 1b).

**Fig. 1.**
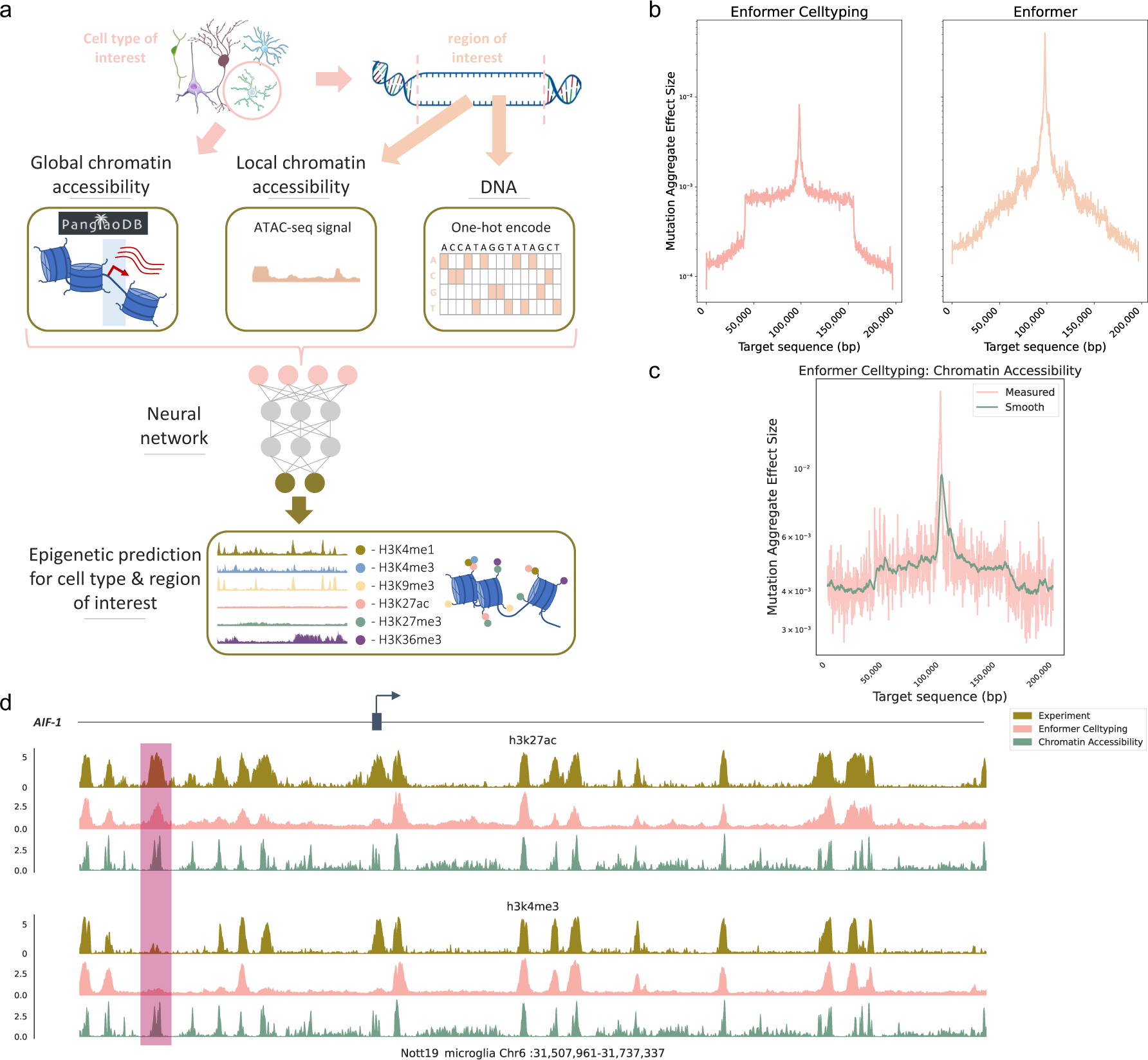
Enformer Celltyping predicts cell type-specific histone marks accounting for distal DNA regulation. (a) Enformer Celltyping uses transformer modules on DNA sequence and embeds both local and genome-wide chromatin accessibility signal to predict cell type-specific histone mark signals. Image created with the help of SciDraw (https://scidraw.io/)^52–55^ (b) The Experimentally derived receptive field for both Enformer and Enformer Celltyping. Random permutations are made at increasing distances from the centre point (100,000 on the x-axis) and the effective change in the prediction on that central point is measured (y-axis). (c) Experimentally derived local chromatin accessibility receptive field for Enformer Celltyping following a similar permutation approach to (b). (d) An example of the observed (Experiment), predicted (Enformer Celltyping) and the chromatin accessibility signal at the transcriptional start site (blue arrow) of the Allograft inflammatory factor 1 (*AIF-1)* marker gene in microglia. The y-axis denotes the −log_10_ p-value, indicating the statistical significance of a protein binding at a genomic position. Highlighted in red is a H3K27ac-specific peak not found in H3K4me3 highlighting that the model uses more than chromatin accessibility signal to identify peaks.

Similarly, we tested the effect of changes in the local chromatin accessibility on Enformer Celltyping by measuring the average change in prediction across reference and alternative chromatin accessibility signal on the centre four positions (512 base-pairs) of the output signal. We replaced the cell type-specific local chromatin accessibility signal of 640 base-pairs with the average signal at increasing distances up to the full receptive field of the model (Fig. 1c). The code for testing the DNA and chromatin accessibility receptive fields are available at: https://github.com/neurogenomics/EnformerCelltyping.

### Visualising cell type embedding

A side-effect of embedding the global chromatin accessibility information in Enformer Celltyping is that these latent representations of cell types is continuously improved by the model during backpropagation. We can then visualise these latent representations with a 2D projection. or all 104 training cell types in our training set we plotted the global chromatin accessibility embeddings using UMAP^48^ with default parameters (Fig. 2a).

**Fig. 2.**
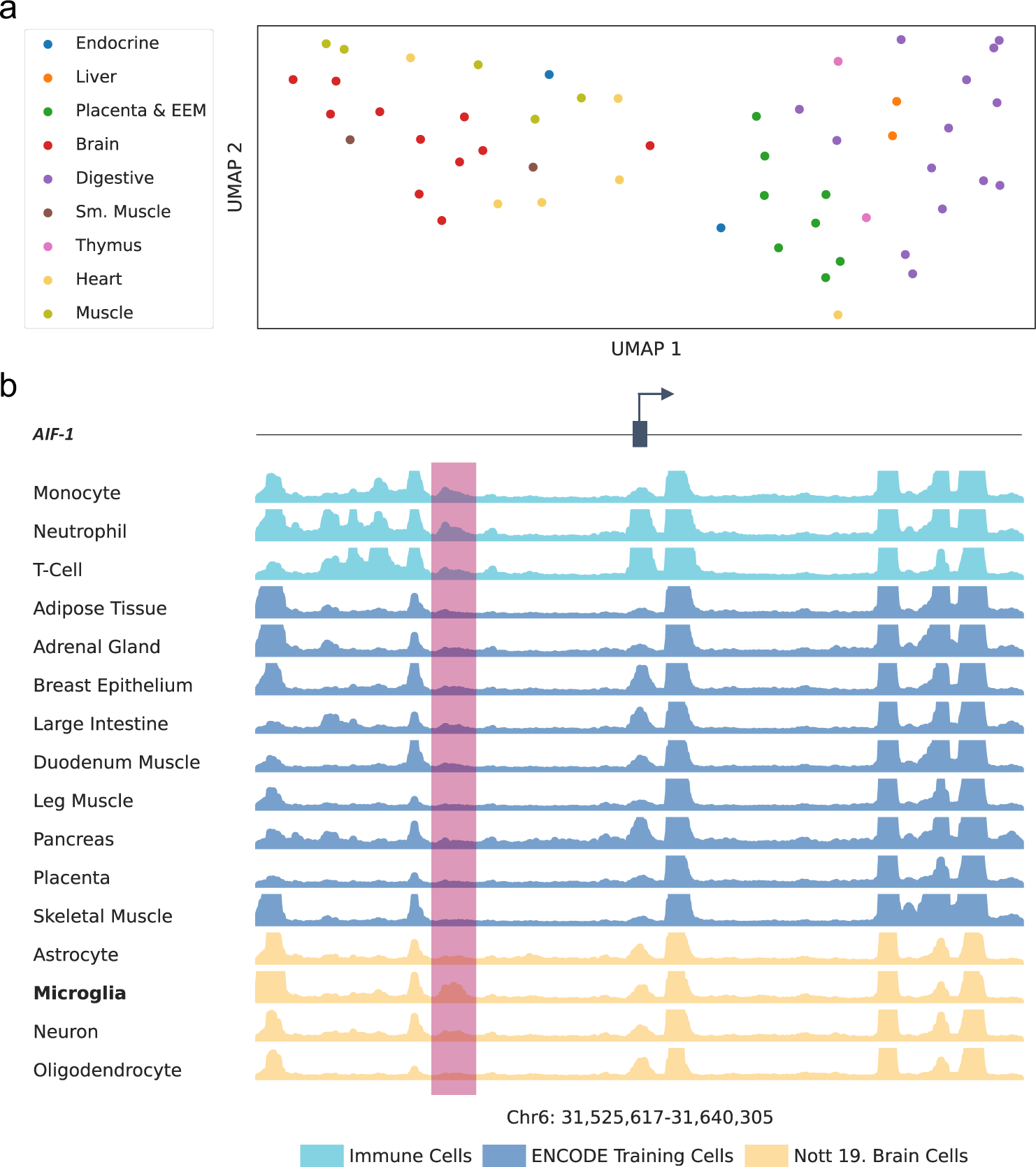
Enformer Celltyping predicts cell type-specific histone marks signals. (a) A UMAP projection of Enformer Celltyping’s global chromatin accessibility embeddings. Cell types from EpiMap are coloured by their broad tissue type. (b) Enformer Celltyping’s H3K27ac predictions at the transcriptional start site of *AIF-1* gene across varying cell types. *AIF-1* is the canonical marker of microglia. Highlighted in red is a microglia-specific peak not found in the other brain cell types but present in immune cells.

### Benchmarking

Enformer Celltyping was benchmarked against Epitome^24^ across the six histone marks for three immune cell types from EpiMap^35^ (CD14+ monocytes, CD16+ neutrophils and naive CD4+T cells) (Fig. 3a). Epitome predicts transcription factor binding and histone marks for a cell type using chromatin accessibility (DNase or ATAC-Seq) data and was shown to have best-in-class predictive performance at cell type-specific histone mark predictions^24^.

**Fig. 3.**
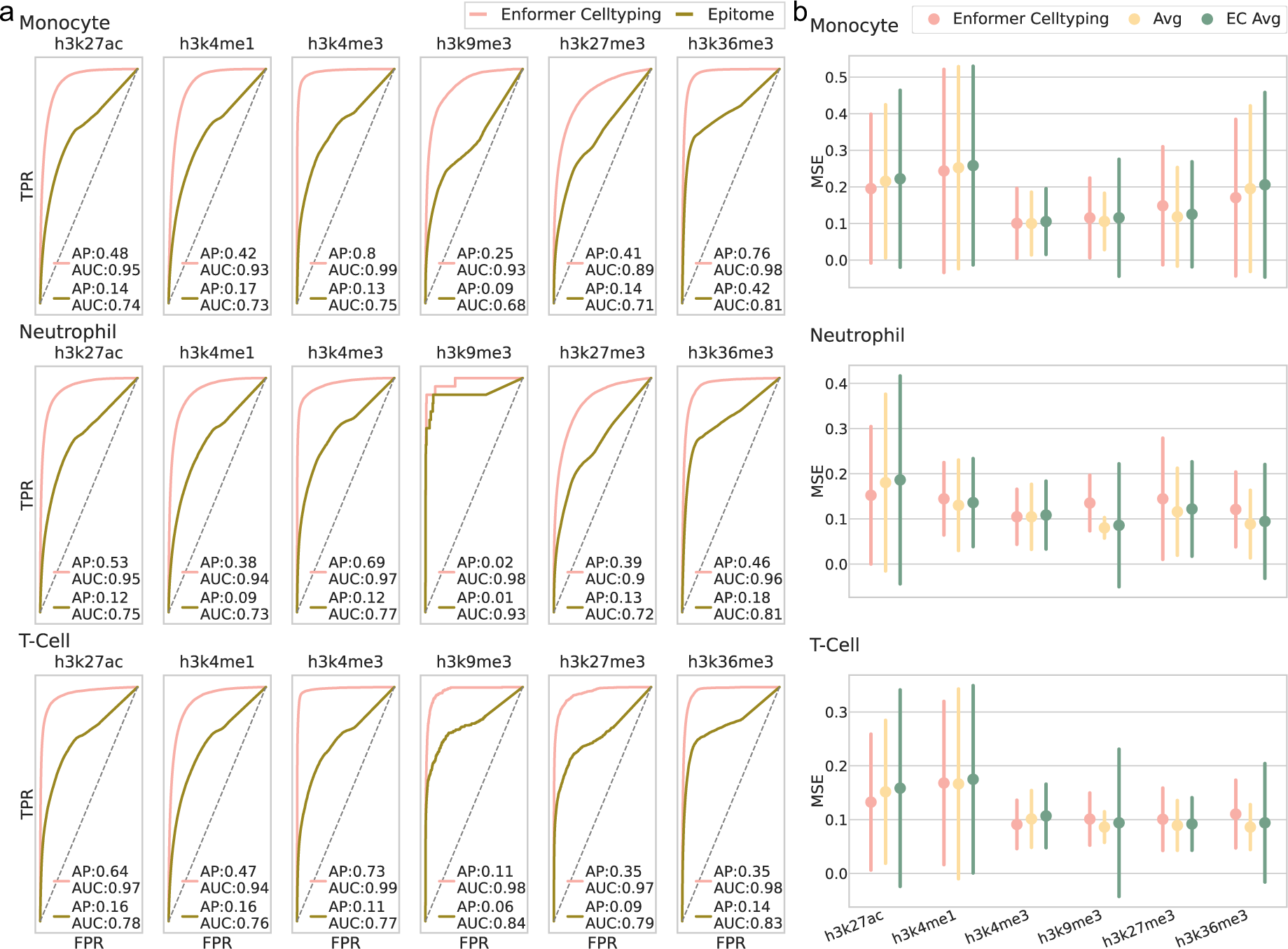
Enformer Celltyping makes best-in-class cell type-specific histone mark predictions. (a) Enformer Celltyping’s outperforms Epitome at genome-wide epigenetic predictions for all three immune cell types (held out from model training). Enformer Celltyping’s predictions were binarized and averaged to 3,200 base-pairs to enable a comparison with Epitome. Note that this binarization and averaging over large genomic regions leads to a blurring of their signals and suboptimal performance overall. The receiver operating characteristics (ROC) curves with the area under ROC score and the average precision (AP) score are reported. (b) Enformer Celltyping frequently, but not consistently, predicts cell type-specific values (for three immune cell types) better than the average signal of the training cell types. The three cell types were not in the training data used to calculate the average. Moreover, Enformer Celltyping’s superior performance against the average predictions from pre-training (EC Avg) highlight the model’s ability to pick up on cell type-specific signals. Performance is measured by the mean squared error (MSE) for genome-wide predictions with error bars for the standard deviation of the error of the predictions genome-wide.

The weights for a pre-trained version of Epitome, trained on Panc1, PC-9, OCI-LY7, MCF-7, Karpas-422, IMR-90, HepG2, HeLa-S3, HCT116, H9, H1, GM23338, GM23248, GM12878, A549 cells from ENCODE^31^, were supplied by the authors and which we have made available through figshare (https://figshare.com/account/home#/projects/159143). Both Epitome and Enformer Celltyping were used to make genome-wide predictions of the same six histone marks for the three immune cell types. Epitome predicts in 200 base-pair bins in a classification setting – whether there is a peak in a 200 base-pair interval or not whereas Enformer Celltyping has been developed to predict signal values directly (-log_10_ p-values) at 128 base-pair resolution. To compare the methods, Enformer Celltyping’s predictions were converted into classification scores using a −log_10_ p-value cut-off of 2, similar to previous work^35^. The predictions for the two models were averaged to the lowest common multiple between the two (3,200 base-pairs). Both of these differences led to a blurring of signal and suboptimal predictions for both approaches but were necessary for comparison. As used in previous work^25^, the performance was evaluated using a balanced measure of performance, accounting for the disparity in the number of regions with and without peaks, the average-precision (AP) for each cell type – histone mark combination. AP is a single score that represents the precision-recall curve based on the rank of predictions using the mean of precisions for each threshold, with the increase in recall from the previous threshold considered as the weight.

### Off-centre correlation analysis

Enformer Celltyping makes histone mark predictions for 128 base-pair bins in a 114,688 base-pairs wide predictive window. We tested whether sliding the genomic location, i.e. the DNA and local chromatin accessibility information, would result in a change in the histone mark prediction at a given genomic location. We tested this by randomly sampling 1,000 genomic positions from our three immune cell types (CD14+ monocytes, CD16+ neutrophils and naive CD4+T cells) which have SNP information in the Blueprint phase 2 hQTL studies data^10^. For each position, we measured the correlation in matching genomic regions between the centred prediction and two off-centre predictions, one up and one down-stream. The positions up and down-stream where chosen to capture the full input window as output (Additional file 1: Fig. S7), to capture as much overlap as possible. Finally we reported the range of these correlations for both genomic data with and without a SNP to ensure genetic variants did not cause a bias in the results (Fig. 6b). The code for testing the off-centre correlation analysis is available at: https://github.com/neurogenomics/EnformerCelltyping.

**Fig. 4.**
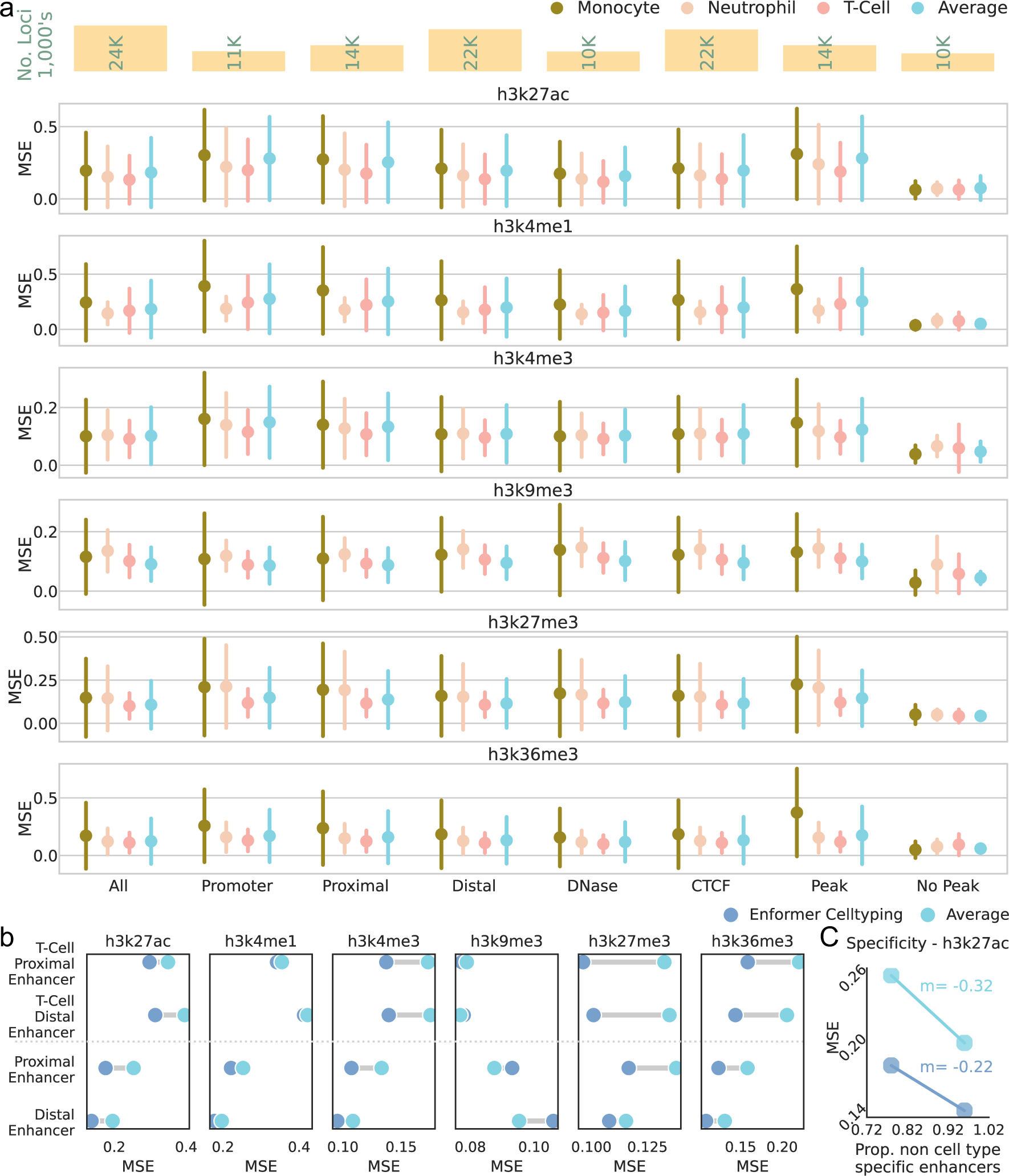
Enformer Celltyping performs better than the training set average at predicting functionally relevant regions. (a) Enformer Celltyping’s (EC) mean squared error (MSE) across different regions sourced from Search Candidate cis-Regulatory Elements by ENCODE (SCREEN)^57^ and derived from a diverse range of ENCODE cell types. Performance is shown against the average signal of the training cell types, for the three immune cell types excluded from the training dataset. “All” represents genome-wide performance and Peak and No Peak regions were derived based on a −log_10_ p-value cut-off of 2^35^. The bars denote the standard deviation of the error of the predictions across all regions within SCREEN that were labelled with the functional annotation. The top panel gives the number of predicted genomic locations (in thousands) containing the functional region, “no peak” count represents a full predicted region (114,688 base-pairs) without any histone mark peaks. (b) Validation by cell type-specific functional annotations highlights EC’s improved performance compared to the training cell type average. EC’s MSE across the two different enhancer genomic regions sourced from SCREEN and derived from a diverse range of ENCODE cell types (“Proximal/Distal Enhancer”) versus derived from T-cells (“T-Cell Proximal/Distal Enhancer”). Performance is aggregated to get the mean of all regions tested and is shown against the average signal of the training cell types. Although the overall performance didn’t always improve, Enformer Celltyping’s performance compared to the average improved (lower MSE) for the vast majority of enhancer types and histone marks by using cell type-specific SCREEN data. (C) For the enhancer regions derived from all cell types, the proportion of non-T-cell-specific loci were identified (y-axis) with MSE performance shown for H3K27ac i.e. the active mark at enhancers (x-axis). The average and EC’s performance improve when there are more regions of no histone activity (higher proportion, lower MSE) but the slope (m) is steeper for the average showing EC’s superior performance in regions with activity.

**Fig. 5.**
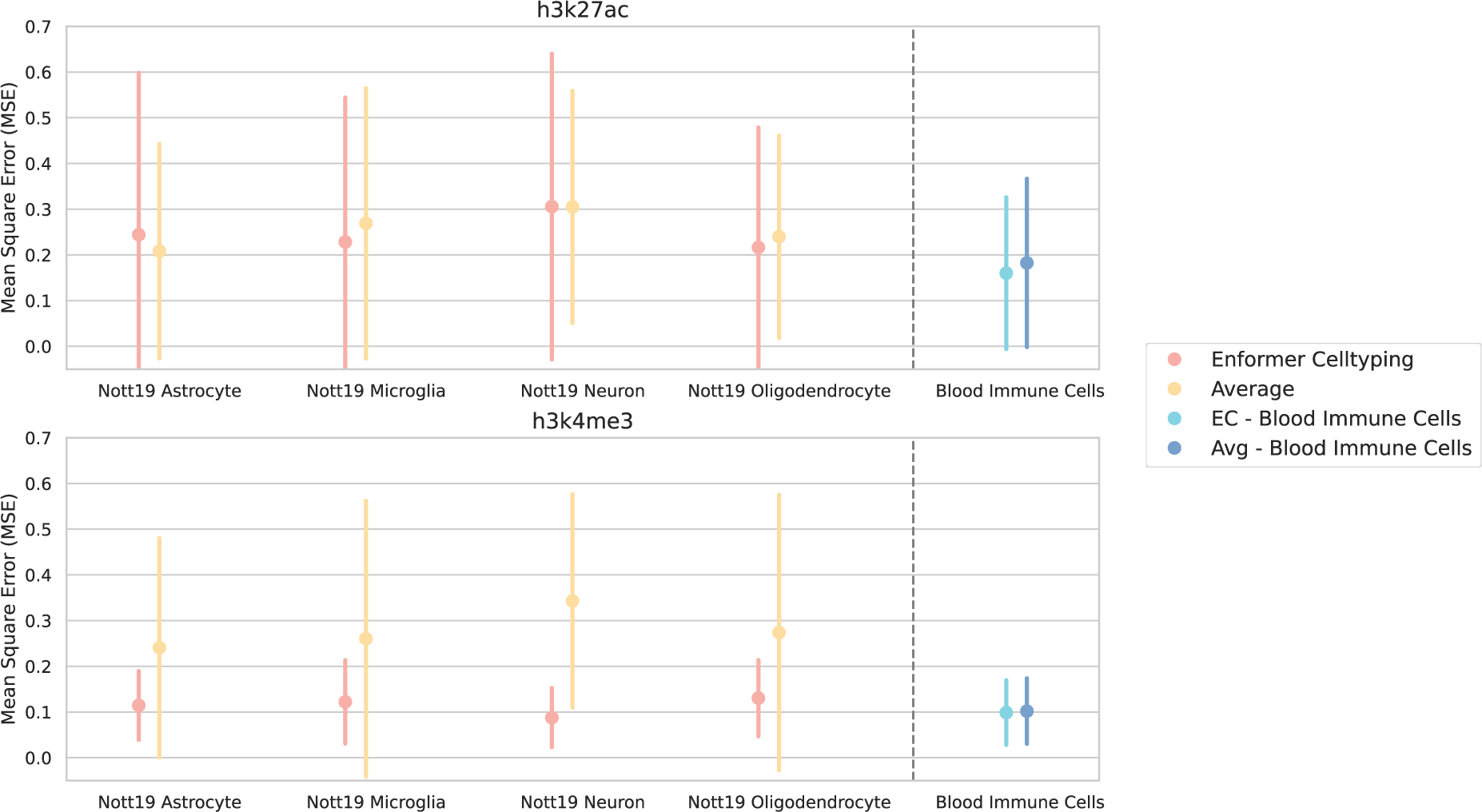
Enformer Celltyping can predict in cell types generated outside of ENCODE. (a) Enformer Celltyping’s mean squared error (MSE) across four isolated cortical brain cell types from Nott *et al.*^40^ and the average signal’s performance. The average performance for Enformer Celltyping and the average signal across the three ENCODE immune cell types (Blood Immune Cells) is also shown for comparison. The results for the “Blood immune cells” shows the union of prediction results for T-cells, Neutrophils and Monocytes. “Average” refers here to the MSE calculated using the average values for the histone mark, across all cell types used in the training of Enformer Celltyping. Performance is measured for H3K27ac and H3K4me3 which were both assayed in the study. Performance is based on genome-wide predictions (over 16,000 per cell type) with error bars for the standard deviation.

**Fig. 6.**
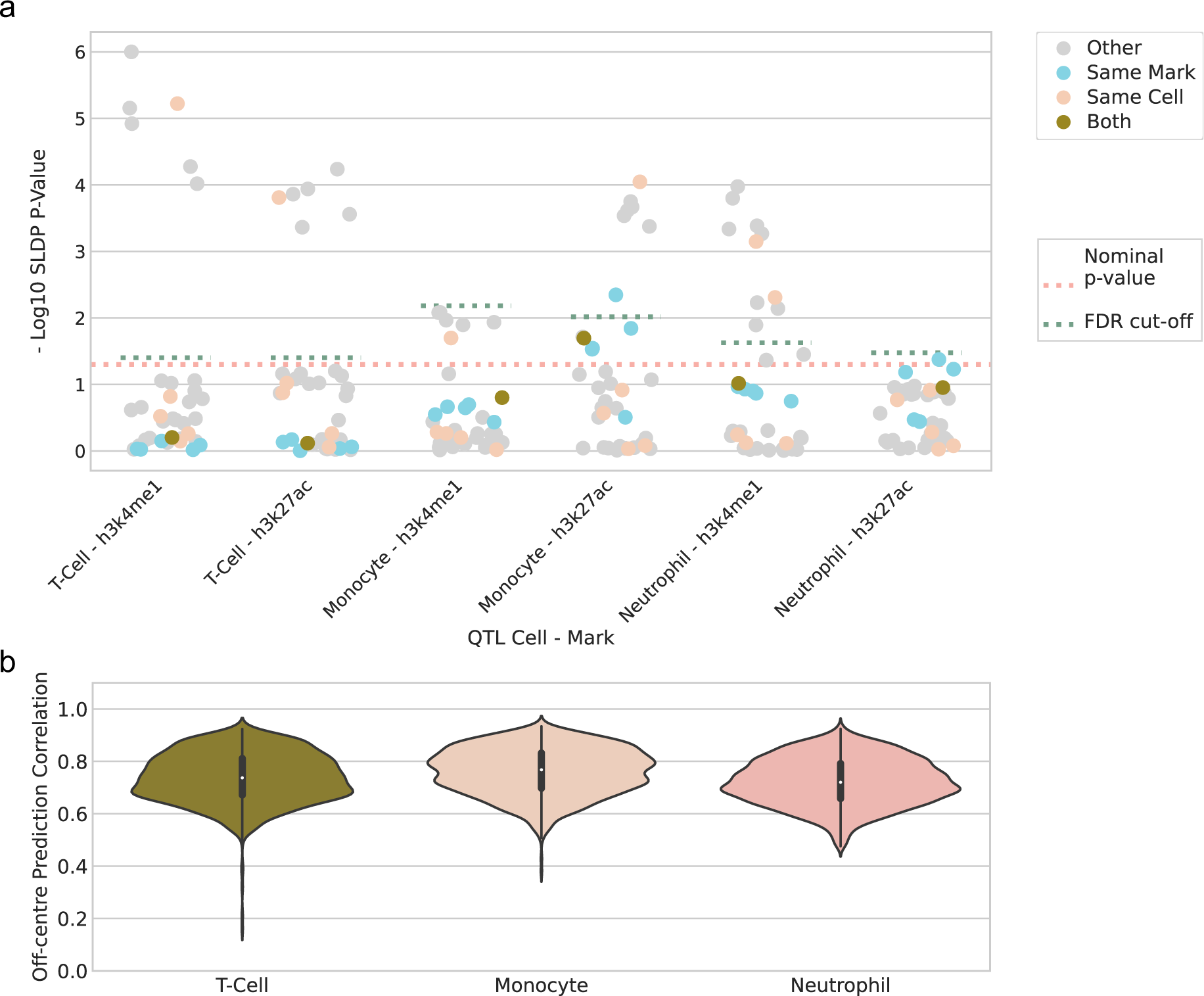
Enformer Celltyping genetic effect predictions predominately do not replicate QTL results. (a) Statistical significance (y-axis) of SLDP^30^ genome-wide concordance between Enformer Celltyping’s genetic variant predictions and measured hQTL effect sizes. The x-axis depicts the six Blueprint phase 2 hQTL studies^10^ and the cell-histone mark predictions are coloured by their relationship to the hQTL study: ‘Both’ – both the histone mark and the cell predicted in were the same as the hQTL study, ‘Same Mark’ – the mark matched, ‘Same Cell’ – the cell matched and ‘Other’ – neither the cell nor mark matched. If there was perfect agreement between our model and the hQTL studies, all ‘Both’ entries would be significant. A nominal p-value (pink dashed line) and the cut-off after Benjamini and Hochberg correction based on the minimum, non-significant p-value, corrected at the level of each hQTL (Turquoise dashed line) are both added. (b) The model makes consistent predictions for a genomic region, regardless of where it lies within the model’s receptive field. The correlation (y-axis) in Enformer Celltyping’s histone mark predictions at matching genomic regions when the model’s input window is moved. This was tested for the three immune cell types (x-axis) for 1,000 genomic positions each with and without genetic variants added.

### Histone Mark QTL SLDP

We predicted the effect of each genetic variant from the six hQTL datasets^10^ with Enformer Celltyping by predicting the histone mark signal for both the reference and the alternative sequence, calculating their difference and summing this effect across the genomic window to get a single, signed score for each SNP. We predicted in six cell types, three matching immune cells and three unrelated cell type from EpiMap: Pancreas, Leg Muscle and Stomach tissue, to act as negative controls. To obtain a prediction across the full input window size, and thus utilising the full receptive field of Enformer Celltyping, we predicted with the SNP first centred in the input window and then performed two further predictions with the SNP slid off-centre (Additional file 1: Fig. S7) and appended these three output windows before calculating the score. Finally, we averaged scores computed using the forward and reverse complement sequence and with small, random sequence shifts up and downstream. This meant for each SNP, we predicted (2x reference and alt, 3x full receptive field, 4 x reverse complement and random shift) 24 predictions. We predicted for 867,568 SNPs across the six hQTL datasets for the six cell types, resulting in approximately 125 million predictions.

We utilised signed linkage disequilibrium profile regression (SLDP)^30^ to measure the statistical concordance between the signed variant effects (our model’s predictions) and the a genome-wide association study’s marginal correlations (the aggregated hQTL SNP values, see Methods section Data collection and processing). SLDP uses generalized least-squares regression to measure the agreement between these, iteratively inverting the direction of the signed variant effect measures along with their neighbouring entries in blocks to derive a null distribution. The method accounts for population linkage disequilibrium (LD) so does not require any fine-mapping strategies. The measured agreement defines how important the variants are to the phenotype’s heritability^30^. We ran SLDP for all combinations of our cell type predictions and histone mark with each hQTL set (6x cell types, 6x histone mark predictions and 6x hQTL sets, resulting in 216 tests) and report the results in Fig. 6a. All code to run the genetic effect preditions and SLDP analysis (with a conda environment) is available at: https://github.com/neurogenomics/EnformerCelltyping.

### Global chromatin accessibility signal – cell type-specific motif enrichment

To inspect the cell type-specific motifs based on the global chromatin accessibility signal to infer what was driving the cell type-specific signal the model captures, we first predicted genome-wide tracks with and without masking the global chromatin accessibility signal using Enformer Celltyping. Histone mark peaks that are dependent on the global signal were identified by checking how much peak strength weakened when the global signal was masked in the set of all peaks from the non-masked predictions (peaks identified as a −log_10_ p-value score >2). For each histone mark, the differences were ordered and the top 10% of peaks reliant on the global signal were identified. The DNA at these positions were run through Homer’s^49^ known motif analysis to search for motif enrichment using default settings apart from setting the size to 128 base-pairs to match Enformer Celltyping’s predictive resolution. To ensure these motifs were specific to the global signal, the bottom 1% of peaks (which gave a similar number of peak regions) for each histone mark were also identified and ran through Homer and any overlapping known motifs were removed from the results (resulted in a median proportion of approximately 10% of significant motifs being removed). The Homer motif analysis is available at: https://github.com/neurogenomics/EnformerCelltyping.

The transcription factors’ genes relating to the resulting motifs were tested for cell type specificity using EWCE^50^ bootstrapping (repeated 10,000 times with a false discovery rate (FDR) adjusted p value threshold of 0.05). All known motifs from Homer were used as the background set (approximately 400 transcription factor genes) after mapping non-human genes to human based on one-to-one orthologs. The single-cell RNA-Seq reference dataset used to determine cell type enrichment was the Descartes human whole body, fetal dataset containing approximately 377,000 cells and 77 distinct cell types^51^. This approach was repeated for all cell types of interest and FDR multiple test correction was implemented to account for the repeated tests (Fig. 7).

**Fig. 7.**
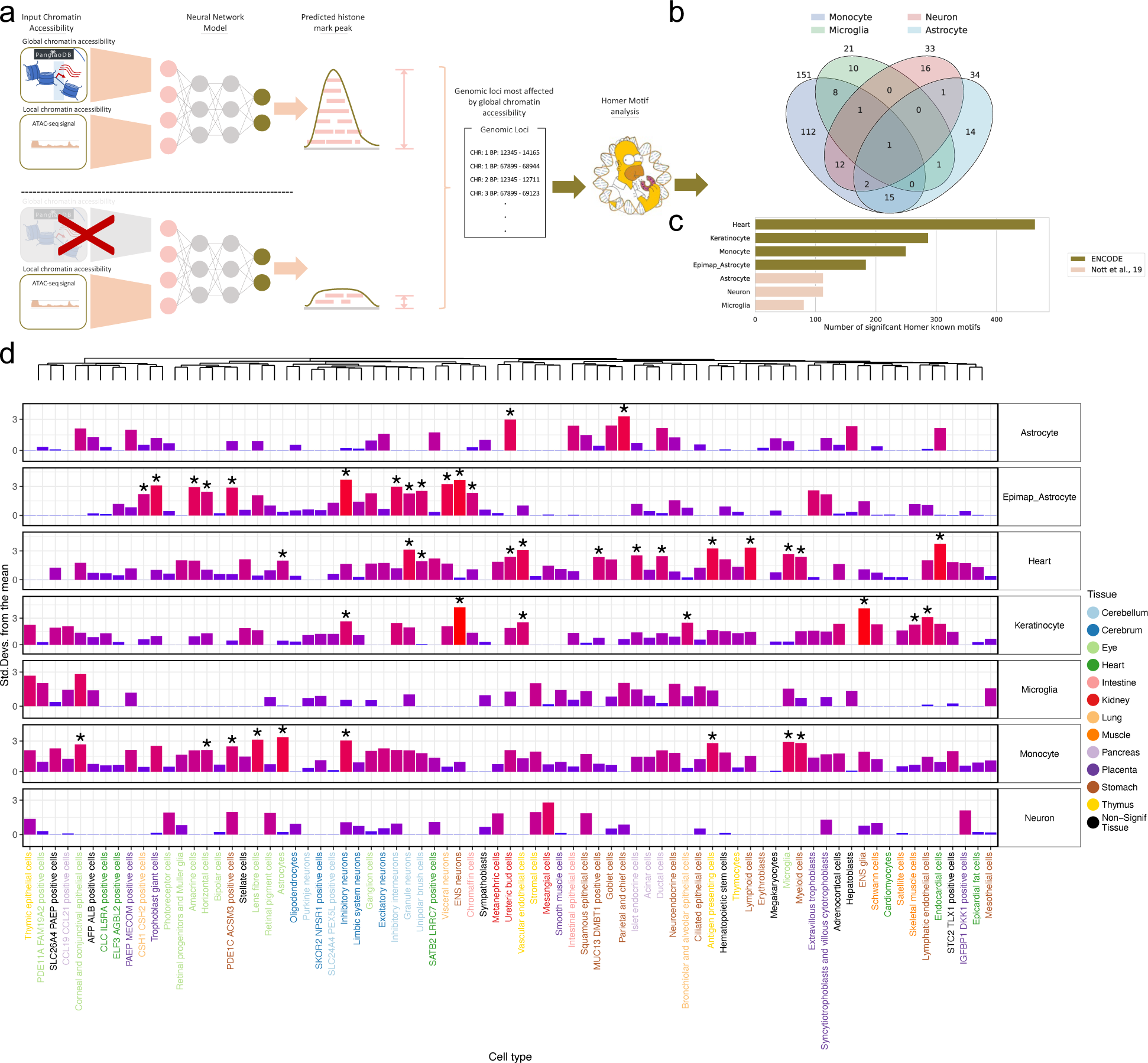
Enformer Celltyping identifies motifs for cell type-specific transcription factors which are enriched when the global chromatin accessibility signal is withheld. (a) Predictions with and without the global chromatin accessibility signal were made to identify peaks and their genomic loci that rely the most on the global input signal. These top loci were analysed with Homer49 for each cell type. (b) The overlap between significant transcription factors across the immune and brain cell types tested for all histone marks. (c) The number of significant transcription factors per cell type. (d) Cell type-specificity of the transcription factors from Homer for each cell type, calculated using EWCE50. The rows are the differ cell types for which predictions were made – note the brain cell types are taken from Nott *et al.*^40^ with the exception of Epimap_Astrocyte which was from ENCODE^31^ sample available through EpiMap^35^. The top row is a dendrogram displaying the clustering of cell types from the Descartes human whole body^51^ single-cell RNA–seq reference dataset used. The columns, ordered by the dendrogram, are the cell types tested from the reference dataset with the colour representing the tissue of origin. Significant associations found by EWCE (false discovery rate adjusted p-value<0.05), indicating enrichment in a cell type are marked with an asterisk. The background geneset were the approximately 400 transcription factor genes from Homer.

## Results

Enformer Celltyping uses Enformer^23^ through a transfer learning approach to expand on its capabilities, predicting in previously unseen cell types. Enformer Celltyping benefits from Enformer’s attention layers which produce its large receptive field (approximately 100k base-pairs), accounting for distal DNA regulatory elements in its predictions (Fig. 1a and Additional file 1: Fig. S1). We measure the sensitivity of this receptive field experimentally with simulated DNA sequences (Fig. 1b). The need for such large receptive field of distal regulatory elements can be inferred by observing the distance between SNPs and the peaks they regulate in histone-QTL studies (Additional file 1: Fig. S2): while these studies are confounded by LD, the prevalence of long-range interactions does not appear to be consistent with a model whereby most histone QTLs act via local motif disruption. Enformer Celltyping can predict in new cell types by embedding global and local chromatin accessibility (ATAC-Seq) signals for the cell type We experimentally show how the model is also sensitive to permutations of the local chromatin accessibility with simulated signals, following a similar trend to the DNA sequence (Fig. 1c). Using this combined approach, Enformer Celltyping shows best-in-class performance at predicting six histone mark signals for any genomic region of interest (Fig. 1d).

Enformer Celltyping embeds a global chromatin accessibility signal for each cell type as well as the local signal. This global cell type embedding shows co-localisation of similar cell types (Fig. 2a). Using this combination of global and local chromatin accessibility signal improved the model’s predictive performance (Additional file 1: Fig. S3), though we note that the local chromatin signal had a larger effect on model performance than the global. Moreover, Enformer Celltyping is capable of making cell type-specific predictions, such as that for microglia at the transcriptional start site of the Allograft inflammatory factor 1 (*AIF-1)* marker gene where a microglia-specific H3K27ac peak was distinct from the predictions for the other brain cell types (Fig. 2b). Enformer Celltyping’s cell type-specific predictive capabilities was notably aided by a pre-training step (Additional file 1: Fig. S4). This step included training the DNA and chromatin accessibility submodules separately to sensibly initialise the weights of the chromatin accessibility layers before combining with the DNA layers which contained the Enformer architecture (see the Methods section Enformer Celltyping architecture).

We benchmarked Enformer Celltyping against Epitome^24^, the current best-in-class approach for predicting cell type-specific histone mark signals, in a genome-wide prediction task on three immune cell types (Fig. 3a). To avoid the performance of our predictions being inflated for this and subsequent benchmarks, we tested three immune cell types (Monocytes, Neutrophils and T-cells) for which data was generated by ENCODE, but which we had held out of the training dataset for Enformer Celltyping. For fair comparison, we binarised the prediction of our model to match that of Epitome (see Methods section Benchmarking). We report the receiver operating characteristics (ROC) curves and the average precision (AP) score for both models. We evaluated performance based on the AP for each cell type and histone mark combination to get a balanced measure of the models’ performances. Enformer Celltyping out-performed Epitome across all cell types and histone mark predictions.

Due to differences in predictive resolution between the two models, large genomic intervals (3,200 base-pairs) had to be used for the classification task so performance could be compared. This would result in suboptimal performance for both models, blurring their signals. To get a clearer picture of how well Enformer Celltyping predicts, following the approach of previous work^5,56^, we benchmarked Enformer Celltyping’s performance against the average signal (Avg) for the same histone mark from the training dataset used to train Enformer Celltyping (Fig. 3b). Here we used the same three immune cell types but benchmarked performance based on the mean squared error (MSE) between the prediction and the actual −log_10_ p-value score, genome-wide at 128 base-pair resolution, utilising Enformer Celltyping’s regression output. We split performance by each histone mark, avoiding reporting a single performance score, to investigate if the model had any specific biases. Overall, Enformer Celltyping performed similarly to the average signal. The predictions for H3K27ac and H3K4me3 signals across the three cell types does consistently out-perform the average, with lower MSE, whereas predictive performance on H3K27me3 consistently under-performs.

One explanation for the inconsistent out-performance relative to the average, is that a model can generally perform well genome-wide by predicting the average signal. To explore this further, we note that during pre-training one of the Enformer Celltyping submodules is explicitly trained to predict the average: it’s performance is shown in Fig. 3 as “EC Avg”. This pre-training step was necessary as Enformer Celltyping uses Enformer as a pre-trained model for the DNA input module. The Enformer-derived layers had sensible weight initialisations whereas the model’s chromatin accessibility module weights were randomly initialised. Thus the short pre-training step avoids issues where all predictions were made based on the DNA information and acts as a warm-up step for the chromatin accessibility layers. During pre-training, the DNA input module predicted the average and distribution of histone mark signals across the training cell types and the chromatin accessibility input module predicted the difference between the average signal and the cell type specific signal (see the Methods section Enformer Celltyping architecture and Additional file 1: Fig. S1a). Subsequent layers combining the two modules were next added for the second, full training step (Additional file 1: Fig. S1b). This pre-training step improved the overall performance of Enformer Celltyping (Additional file 1: Fig. S4), as did including the distribution of histone mark signals as well as the average in the DNA module predictions (Additional file 1: Fig. S5).

Enformer Celltyping’s cell type-specific predictions (Enformer Celltyping) outperform that of the average predictions from pre-training (EC Avg) in the vast majority of cell types and histone marks. Moreover the variance in the error of the predictions genome-wide is far smaller, indicating how Enformer Celltyping picks up on cell type-specific signals even at genome-wide predictive performance comparisons (Fig. 3b).

While these results show Enformer Celltyping performs similar to the average signal, this is genome-wide performance, something that the average signal would be expected perform strongly at given the overlap between cell’s histone mark signals. To investigate whether Enformer Celltyping performs better than the average at predicting functionally relevant regions (e.g. peaks, enhancers and promoters), we measured the model’s predictive performance across a number of genomic region types (Fig. 4a). These focused on differences in the histone mark signal (Peak and No Peak regions based on a −log_10_ p-value cut-off of 2^35^) and on different biological regions from Search Candidate cis-Regulatory Elements by ENCODE (SCREEN)^57^, available from: https://screen.encodeproject.org/. Overall, Enformer

Celltyping performs better (that is, has a lower MSE) in known functional genomic regions whereas the model performs slightly worse (relative to the average’s MSE) in regions of no histone mark activity (No Peak). For example, in Fig. 3b, it can be seen that predictions for H3K27me3 in T-cells had higher MSE (worse) than the average signal genome-wide, however, in Fig. 4a, it performs better in all functional regions but worse in “no peak” regions. This could explain how genome-wide, Enformer Celltyping does not out-perform the average given the prevalence of genomic regions with no histone mark activity. The model’s performance is notably strong for distal enhancers genomic regions for the active enhancer marks - H3K27ac^58^ and H3K4me1^59^, given enhancers’ cell type-specificity^60^. However, it is worth noting that Enformer Celltyping performance varies by cell type and similar to our findings in Fig. 3b, the model performs worse in monocytes with higher variance in the error of the predictions with an exception being in regions without histone mark activity (No Peak): an explanation for this could be that the models predicts fewer peaks for monocytes, and thus performs better at predicting regions without peaks, but worse at all regions which should have peaks.

It should be further noted that this point has been made using a definition of functional regions from across all cell types: the advantage in performance of Enformer Celltyping may be greater still if cell type-specific definitions of functional regions were used. To test this, we sourced the SCREEN functional enhancer regions based solely on T-cells, available from: https://screen.encodeproject.org/, and tested performance of the training set average and Enformer Celltyping’s T-cell predictions on this subset of regions (Fig. 4b). Although the overall performance didn’t always improve, performance compared to the average improved (lower MSE) for the vast majority of enhancer type and histone mark combinations by using cell type-specific SCREEN data with the largest improvements noted in the distal repressive marks (H3K27me3 and H3K36me3). This trend highlighted how genomic deep learning models will perform better in regions of no activity which was likely added to by the use of MSE to measure performance. We investigated this relationship further by calculating the proportion of non-T-cell-specific loci in the SCREEN enhancer sets and evaluated the relationship against model performance in T-cells on H3K27ac; the canonical active enhancer mark (Fig. 4b). The training set average and Enformer Celltyping’s performance improved when there are more regions of no histone activity in T-cells (higher proportion, lower MSE) highlighting this trend. However, the slope is steeper for the average, showing Enformer Celltyping performs better in cell type-specific functional regions like enhancers.

Up to this point, Enformer Celltyping’s performance has only been tested on cell types generated from ENCODE. We next measured its performance in a manner that replicates how researchers can use Enformer Celltyping to predict histone marks in their cell types of interest^56^. To this end, we utilised H3K27ac and H3K4me3 ChIP-Seq and ATAC-Seq data for isolated cell types (PU.1+ microglia, NeuN+ neuronal, OLIG2+ oligodendrocyte and NeuN-LHX2+ astrocytes) from resected cortical brain tissue^40^. Consensus peak p-value tracks (taking individuals as replicates) were derived and the data processed in the same manner as for the training cell types. Enformer Celltyping has not been trained on any isolated cell types from the brain and this data was not collected following the ENCODE protocols so its performance across the two measured histone marks should give a fair indication for any future applications of the model (Fig. 5). To enable a fair comparison between ENCODE and non-ENCODE cell types, we again tested three ENCODE immune cell types which were held out of the Enformer Celltyping training dataset: here their results have been grouped and they are referred to as “Blood immune cells”. Overall, Enfomer Celltyping’s performance did not worsen drastically (relative to the ENCODE blood immune cells) despite the cells’ dissimilarity to ENCODE samples. This becomes more impressive when we consider the performance of the average signal which showed a sharp increase in the MSE for H3K4me3 in the non-ENCODE cell types, relative to the ENCODE blood immune cells. Enformer Celltyping’s ability to predict well in the presence of such a domain shift compared to the average signal of the ENCODE training cell types, highlights its ability to use the cell type-specific chromatin accessibility effectively when making predictions.

Having validated Enformer Celltyping’s ability to predict the cell type-specific, genome-wide histone mark signals conditioned with chromatin accessibility information, we proceeded to test its primary application of predicting the cell type-specific effect of non-coding, disease-relevant genetic variants on the epigenome. The requirements of this task are why Enformer Celltyping was designed in the manner that it was: able to predict in novel cell types (such that it can predict SNP effects in disease relevant cells); as well as incorporating the effect of distal genetic regulators (which we showed to be prevalent in Additional File: Fig. S2).

We developed an approach, similar to previous studies utilising QTL datasets^23,43,30^, systematically measuring the genome-wide correlation between our model’s predictions and the measured effects from the QTL studies using SLDP^30^ (see Methods section Histone Mark QTL SLDP and Data collection and processing). We used QTL datasets as they directly measure the effect of extant human genetic variation. Furthermore, we have not concentrated on predicting the effect of genetic variants in isolation due to the confounding effect of linkage disequilibrium (LD), obscuring the coinherited and casual SNPs^30^. Approaches like fine-mapping could help identify the casual SNPs, however recent works have highlighted that despite considerable advancements there is still opportunity for improvement^20,21,22^. Note that past work concentrated on using eQTL studies and predicting genetic variant effects in the same cell types that the models were trained on. On the other hand, we utilised hQTL datasets and our cell type-agnostic approach to match the studied cell type, giving insight into the model’s predictive ability of SNP effects on the epigenome. Moreover, we implemented strict pre-processing criteria, filtering out any interactions from the hQTL set where the distance was greater than the model’s receptive field and measured the effect across the full receptive field of the model.

We used hQTL data from the Blueprint project phase 2^10^, for two histone marks (H3K4me1 and H3K27ac), each of which was obtained for three cell types (T-cells, Monocytes and Neutrophils). These were the same cell types which were held out of the Enformer Celltyping training data, as discussed above. These hQTL datasets were generated using whole genome sequencing from neutrophils (n = 196), monocytes (n = 194) and T-cells (n = 169) individuals. We note that while these are the largest published hQTL datasets, they remain small by the standards of modern eQTL datasets, and so it can be considered underpowered to detect QTL effects. We tested 867,568 genetic effect predictions for all of Enformer Celltyping’s predicted histone marks for six cell types against the six hQTL datasets^10^. Similarly to model training, we tested the effect on the forward, reverse strand and with small random shifts in genomic starting position to return the average effect. We expected to see a significant result for the prediction in the same cell type and histone mark for which the hQTL study was completed (Fig. 6a).

The results of the SLDP genetic effect predictions were worse than what was expected with only one of six of the matching histone mark and cell types being nominally significant and none after false discovery rate (FDR) correction of 0.05 (Fig. 6a). We noted that one possible causes of differences, relates not to the quality of SNP effect predictions, but to differences in predictions of peaks for the reference sequence: sometimes Enformer Celltyping doesn’t predict peaks which are seen in the experimental data. If a peak is missed in the predictions, then the effect of a SNP which decreases the size of that (missing) peak, cannot be captured. To assess whether this affects the results, we tried filtering out SNP-to-histone QTLs associated with peaks that were missed by Enformer Celltyping, and re-running the SLDP analysis. However, this did not improve the significant matched cell type and mark cases (Additional file 1: Fig. S6).

In making these predictions, we altered our approach to *in silico* mutagenesis, relative to the standard in the field (Additional file 1: Fig. S7). Our issue with the standard approach, which centres the SNP in the input window, is that it does not utilise the full receptive field of these models (Additional file 1: Fig. S7a-b). Our approach captures the full receptive field by making multiple predictions for each SNP, sliding the input window (Additional file 1: Fig. S7c). Given our approach results in multiple predictions of the same genomic locations, we sought to confirm that predictions were consistent for the same genomic region, regardless of where it appears in the input window (Fig. 6b) (also see Methods section Off-centre correlation analysis). This resulted in extremely high correlations for all tested cell types, confirming that centring the model’s predictions on the SNP and shifting the input window in the hQTL dataset would not affect the histone mark binding predictions.

We have demonstrated that Enformer Celltyping accounts for changes in local chromatin accessibility for its predictions (Fig. 1c) but also that it considers global, cell type-specific signals (Fig. 2, Fig. 5). To investigate what was driving the model’s cell type-specificity, we identified the genomic regions with histone mark peaks that had the highest contribution from the global chromatin accessibility signal for a selection of cell types (Fig.7a and Methods section Global chromatin accessibility signal – cell type-specific motif enrichment). The corresponding DNA sequences for these peaks were assessed for enrichment of known motifs using Homer^49^ (Fig.7b-c). Finally for each cell type, based on the set of significant known motifs, we looked for cell type specificity of the related transcription factors using EWCE^50^ (Fig. 7d).

Overall, Fig. 7 highlights how Enformer Celltyping identifies DNA motifs relating to distinct transcription factors for the different cell types tested and some of these transcription factors match the same cell types in their transcriptional specificity. For example, monocytes had significant enrichments in multiple macrophage and myeloid cell types (Microglia and Myeliod cells and Antigen presenting cells). Furthermore, the strongest enrichment for heart is endocardial cells. Neurons had no significant cell type associations but do show some enrichment in neuronal cell types. Keratinocytes (a subtype of epithelial cells) had significant enrichments in epithelial cell types (Bronchiolar cells and alveolar epithelial cells). Despite this, keratinocytes did also have enrichment in multiple endothelial cells, the meaning for which is less clear. The differentiated astrocyte cell from ENCODE^31^ (Epimap_Astrocyte) showed significant cell type enrichments in brain cell types, predominately in neurons but not in the reference astrocyte cell type. Moreover, we see overlap in the transcription factors identified with Homer in similar cell types such as across the immune cells; microglia and monocytes (Fig.7b). Interestingly, the cell types from Nott *et al.*^40^ (Astrocyte, Microglia and Neurons) showed fewer cell type enrichments than for those from ENCODE (see Epimap_Astrocyte vs Astrocyte for a comparison of similar cell types). We can also see that these cell types found fewer transcription factors from the global chromatin dependent peak motifs (Fig. 7c). This could indicate a bias in Enformer Celltyping whereby, although the domain shift of predicting in isolated, single cell types outside of ENCODE does not seem to affect the genome-wide predictive performance, it does affect the global cell type-identity the model learns. We can also see this ffect when we consider the projection of the model’s embedded global chromatin accessibility signal for the Nott *et al.* brain cell types versus the ENCODE brain cell types which seem to separate in a lower dimensional space (Additional file 1: Fig. S8).

## Discussion

We report a cell type-agnostic model to predict epigenetic signals with the largest receptive field to date of 100,000 base-pairs. Enformer Celltyping achieves this through a transfer learning approach based on Enformer, a model which predicts gene expression and epigenetic signals limited to the same cell types it was trained on^23^. Here, we recreated Enformer, removing the output and convolutional layers after the transformer block and freezing the weights in pre-trained layers. This approach differs from past applications which mostly fit a linear model on top of Enformer’s output^27,28^, offering more flexibility and aligning with current approaches in the machine learning domain such as NLP^61^.

Our approach shows the power of transfer learning using large, pre-trained models in computational biology. Here, we trained Enformer Celltyping on a data set several times larger than Enformer’s, using a fraction of computational resources (132 versus 5,376 GPU hours). We believe this approach, which we have made openly available, shows promise for what researchers can achieve fine-tuning large, pre-trained models in the face of limited resources.

Enformer Celltyping expands on the Enformer model by including chromatin accessibility data to predict in previously unseen cell types. Enformer Celltyping’s architecture is split into two submodules, the DNA module predicting the average/distribution of the histone mark signal and the celltyping module which predicts the difference between the average and the cell type-specific signal. The submodules were trained separately for a short, pre-training step before being combined with subsequent layers to make cell type-specific predictions, ensuring both submodule influenced Enformer Celltyping’s predictions at a given genomic position and improving performance overall (Additional file 1: Fig. S4). The celltyping submodule is, itself, made up of two parts; a global and local chromatin accessibility embedding (adding both of which improved performance Additional file 1: Fig. S3). The global signal, derived from the transcriptional start site of marker genes from PangloaDB^45^, enables the model to encode cell type-specific differences independent of the genomic location. This cell type embedding extends to new cell types, something we demonstrated on cell types collected both as part of and separately to ENCODE^31^ and EpiMap^35^. One critique of our method, evident in Fig. 3b, is that the performance of Enformer Celltyping was not consistently stronger than the average prediction model (EC Avg). This was, to a certain extent, expected as inferring the cell type-specific differences is a harder task than deriving the average signal which is dependent on the quality of the chromatin accessibility data. Here, due to a lack of availability, we used imputed chromatin accessibility ATAC-Seq data for the majority of cell types from EpiMap^35^. A possible improvement could be to use assayed chromatin accessibility data our use a better performing imputation technique^25,62^. Our reliance here on imputed data, highlights the importance of generating new biological datasets specifically designed for training machine learning models: At present almost all models in this field use ENCODE, which was not created for this purpose.

Enformer Celltyping shows strong predictive performance of histone marks in all tested cell types and biological regions. We benchmarked the model against Epitome^24^, the current best-in-class model for cell type-agnostic histone mark predictions, which it out-performed in all histone mark and cell type combinations. There were discrepancies between the model which caused challenges for the benchmark – classification vs regression outputs and differences in bin size. Enformer Celltyping was developed as a quantitative model, predicting a continuous −log_10_ p-value signal, which has been shown to yield better generalisation and interpretability than binary, classification models^63^. The −log10 p-value derived after MACS 2.0 peak calling was chosen over the fold change measure as it typically has a higher signal-to-noise ratio^25^. Moreover, Enformer Celltyping showed strong genome-wide predictions even with a domain shift to isolated cell types assayed outside of ENCODE (Fig. 5). However, this distributional shift did seem to affect the global cell type embedded signal for these cell types (Additional file 1: Fig. S8).

One drawback of Enformer Celltyping’s predictive approach is that predictions are made in 128 base-pair bins. This is lower resolution than the standard bin size after peak calling from ENCODE^31^ (25 base-pairs). A major goal of developing Enformer Celltyping was to use it to predict the effect of genetic variants. Doing so at a lower resolution will dilute the effective change on the epigenetic signal which may have contributed to the suboptimal performance of Enformer Celltyping on the genetic variant analysis. Predicting in 25 base-pair bins was not possible with our transfer learning approach using the frozen layers from Enformer. Possible solutions to enable this would be to either allow the weights to update in Enformer so it can adjust to the desired resolution which would incur a much higher a computational cost, equivalent to that which the original authors faced or to use a U-Net architecture on top of Enformer to increase the resolution, as proposed recently^64^.

We show that Enformer Celltyping is sensitive to changes in DNA sequence, as well as local and global chromatin accessibility information. Both Enformer Celltyping and Enformer’s average effect of genetic variants decreases as distance increases, something which has been independently confirmed by Karollus *et al.*^28^. Interestingly, there is a sharp decrease in the average effect size for Enformer Celltyping after 50,000 base-pairs which was not apparent for Enformer, and may be due to the interaction between the chromatin accessibility and DNA modules. Upon investigating the causes of the model’s predictions based on global cell type information, we found that cell-type specific histone mark predictions are influenced by the presence of cell type-specific transcription factor motifs. However, a number of non-specific cell type enrichments remain, which could be due to confounders such as the limited number of transcription factors with known motifs. Moreover, mapping these back to human based on orthologs and the fact that some motifs will span whole transcription factor families may affect the quality of the results.

A primary goal of machine learning models that predict gene expression or epigenetic signals from DNA, including Enformer Celltyping, is to predict the effect of genetic variants^23,43,30^. Usually, these models can only make predictions in the same cell types they were trained in. We have expanded this utility to a cell type-agnostic approach, enabling genetic variant predictions in any cell type of interest.

Our approach to *in silico* mutagenesis built on that used by Enformer. It matched Enformer’s approach to predict the forward and reverse strand and also with a small random shift in DNA, however we differed on two main fronts: Firstly, we predicted the genetic effect across the full receptive field of the model (Additional file 1: Fig. S7), whereas to the best of our knowledge, only a single prediction was made for Enformer with the SNP centred. Secondly, we had to make separate predictions for each cell type of interest because of our cell type-agnostic approach, whereas for Enformer, only one prediction pass is needed. We ran predictions for 867,568 SNPs from six hQTL studies in six cell types. This totalled approximately 125 million predictions, a significantly more computationally intensive task than that for Enformer (because their one output head covered all cell types and assays).

We performed the first evaluation on *in silico* mutagenesis performance using hQTL data in combination with SLDP^30^. Previous studies have evaluated performance using eQTL data, which is several layers further removed from the presumed direct biological effect of SNPs (eQTLs aggregate over multiple layers of gene regulation, while hQTLs will be more directly affected). Moreover, other studies have used fine-mapping to evaluate model performance, but such approaches are compromised by their reliance on fine-mapping approaches, which differ depending on the methodology used^20,21,22^. When previous studies have used SLDP with eQTL data, they have also only considered the z-score as opposed to the actual p-values computed^23^. We believe this framework for testing models provides an objective standard for performance based on p-value cut-offs.

We have sought to define a more structured framework for SLDP evaluation of QTLs, including utilising the full receptive field of the model in question (Additional file 1: Fig. S7). As part of this, we wanted to ensure that centring on the genetic variant of interest rather than the affected epigenetic region (or for other approaches gene expression) would not affect the prediction results (Fig. 6b). Strong correlations show that the model predicts similar signals regardless of where the genomic region is positioned in the input window. We also filtered out any of the hQTL SNP to histone mark binding sites that were outside of the receptive field of our model (∼100,000 base-pairs)This important step was omitted from previous approaches^30^.

Overall, Enformer Celltyping’s performance in predicting the effect of genetic variants predominately did not replicate the QTL results using the SLDP measure of concordance^30^ (Fig 6a). We hypothesised that two structural issues with the design of the model contribute to this, while also noting that the training and validation data probably need to be improved as well. Firstly, that the hQTL set contained interactions which caused a histone mark binding position to be removed by a genetic variant but these histone mark binding positions were not captured in Enformer Celltyping’s prediction on the major allele. Secondly, that Enformer Celltyping struggles to correctly predict the effect of genetic variants. We tested the effect of the first issue by removing such interactions before running SLDP but this had little effect on increasing the number of expected significant associations (Additional file 1: Fig. S6). The second issue, which received growing attention in the literature, is specifically related to Enformer, and due to transfer learning, also affects Enformer Celltyping. Firstly, Karollus *et al.*,^28^ showed how the predicted effect of regulatory elements decreased with distance and did not match the measured causal effects of these elements, such as enhancers. This means that on average, Enformer struggles to identify distal regulatory genomic regions, weighting their importance less than local regulatory information. This may give insight into Enformer Celltyping’s relatively poor performance in predicting the effect of genetic variants where the majority of genetic variants tested were distal (Additional file 1: Fig. S2). Secondly, Sasse *et al.*,^27^ comprehensively showed how Enformer underperforms at predicting the effect of genetic variants, even predicting the incorrect direction of effect in up to 40% of tested cases. The authors hypothesised that this was a result of Enformer being trained on points across the reference genome and then needing to make out-of-sample predictions for individuals’ personal genetic variants at any locus. Alternative methods use an across-individual approach, such as PrediXcan^65^ which was trained directly on the genetic variance across samples. The authors go on to show how the genomic regions Enformer uses when making predictions do not overlap with the loci containing the genetic variants. Approaches like PrediXcan force the model to consider loci with genetic variants by directly training on these. Both these analyses were conducted on Enformer’s expression predictions but we believe they also hold for the predicted epigenetic marks of interest in our work. These issues are compounded by the fact that models like Enformer Celltyping, which try to predict the effect of genetic variants in previously unseen cell types, need alternative genomic information to infer the cell type it is predicting in. For Enformer Celltyping, this is chromatin accessibility information. However, these experimental datasets generally do not account for the genetic variant thus further disadvantaging the model’s ability to predict the genetic effect accurately. To summarise, we agree with Sasse *et al.*,^27^ that future work hoping to use a machine learning model to predict the effect of genetic variants should concentrate on training models on data which accounts for genetic variants and secondly, that research should focus on predicting in the same cell types upon which the model was trained rather than extrapolating to previously unseen cell types using experimental datasets. This is, of course, limited by the current lack of suitable public data, specifically for epigenetics.

The most substantial future improvements in machine learning for genomics will be led by the data that is available to train and validate the models. Our ability to validate model performance at predicting the effect of genetic variants is limited by our experimental knowledge of variant effects. The hQTL datasets that we have used here are the largest available: They were generated using only ∼196 individuals, which is low for QTL studies and dramatically underpowered by commonly accepted standards in genetics i.e. GWAS. The training data is also limited because datasets pairing whole-genome sequencing and epigenetic data from the same individuals is seldom available. As a result, training assumes that the epigenetic data is linked to the reference human genome, and thus ignores the effect of variation which must exist. Advancement in the field should be based on generation of a focused dataset where these problems are resolved for at least one cell type.

In summary, our development of Enformer Celltyping, with its ability to predict in novel cell types, made it possible to evaluate whether *in silico* mutagenesis accurately captures experimentally measured effects of genetic variants on the epigenome. We have found that despite having the capacity to make accurate, genome-wide epigenetic predictions in previously unseen cell types, Enformer Celltyping like other genomic deep learning models, can not yet accurately predict the effect of genetic variants.

## Conclusion

Enformer Celltyping is a self-attention based, neural network model which predicts histone mark activity in previously unseen cell types. It makes these predictions based on DNA sequence and chromatin accessibility information. Our model uses a transfer learning approach on a pre-trained Enformer^23^ model to account for the distal genetic regulatory effects and embeds both a global and local representation of cell types from chromatin accessibility information. We show Enformer Celltyping’s predictions out-perform current approaches and generalises to cell types assayed separately to the original training set. However Enformer Celltyping, like other similar models, can not predict the effect of genetic variants adequately yet. We have made a pre-trained version of Enformer Celltyping available along with our transfer learning approach.

## Data availability

All training and test cell types from EpiMap, along with scripts to download and complete all pre-processing steps are available at https://github.com/neurogenomics/EnformerCelltyping. Blueprint phase 2, histone mark QTL data was downloaded from the European Genome-Phenome Archive (ID: EGAD00001005199). The full training and all validation scripts are also available at https://github.com/neurogenomics/EnformerCelltyping. A list of the training regions and cell types, the one-hot encoded DNA sequence, the trained Enformer Celltyping model’s weights, the average chromatin accessibility signal of the training cell types are all available through figshare (https://figshare.com/account/home#/projects/159143).

## Code availability

The Enformer Celltyping model architecture, training scripts and all analysis are available from https://github.com/neurogenomics/EnformerCelltyping.

## Acknowledgements

I would like to thank Leo Linden, Simon Mathis, David Kelley and Jacob Schreiber for their invaluable conversations and feedback that supported this work. I would also like to thank Alyssa Morrow for her help supplying a version of Epitome for benchmarking. This study makes use of data generated by the Blueprint Consortium. A full list of the investigators who contributed to the generation of the data is available from www.blueprint-epigenome.eu. Funding for the project was provided by the European Union’s Seventh Framework Programme (FP7/2007-2013) under grant agreement no 282510 – BLUEPRINT.

