## Additional file 1 for "Predicting cell type-specific epigenomic profiles accounting for distal genetic effects"

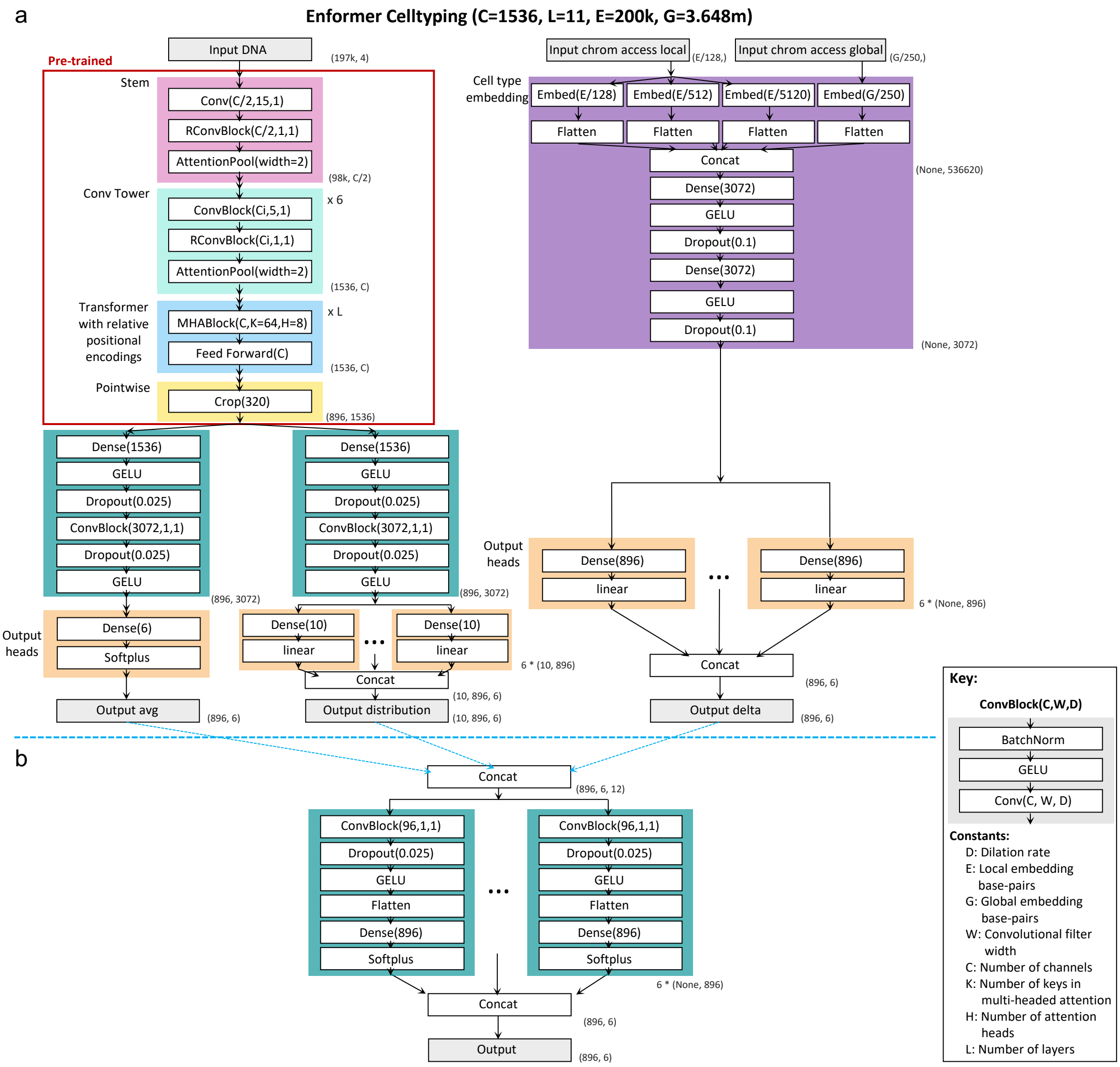


**Fig. S1 Enformer Celltyping architecture**. (a) pre-training, warm-up step: Enformer Celltyping's architecture is split into two separate submodules – The DNA half (stemming from 'Input DNA') which inputs DNA sequence and uses a pre-trained, chopped, frozen version of Enformer through a transfer learning approach to predict an average histone mark signal ('Output avg') and distribution ('Output distribution') across training cell types for the genomic region; and the celltyping half (stemming from 'Input chrom access local' and 'Input chrom access global') which embeds chromatin accessibility data from the same region and predicts the difference between the average histone mark signal and the cell type-specific signal ('Output delta'). (b) Second training step: Enformer Celltyping combines the three outputs

(‘Output avg’,‘Output distribution’ and ‘Output delta’) with subsequent layers to predict the cell type-specific histone mark signal for a given genomic region.

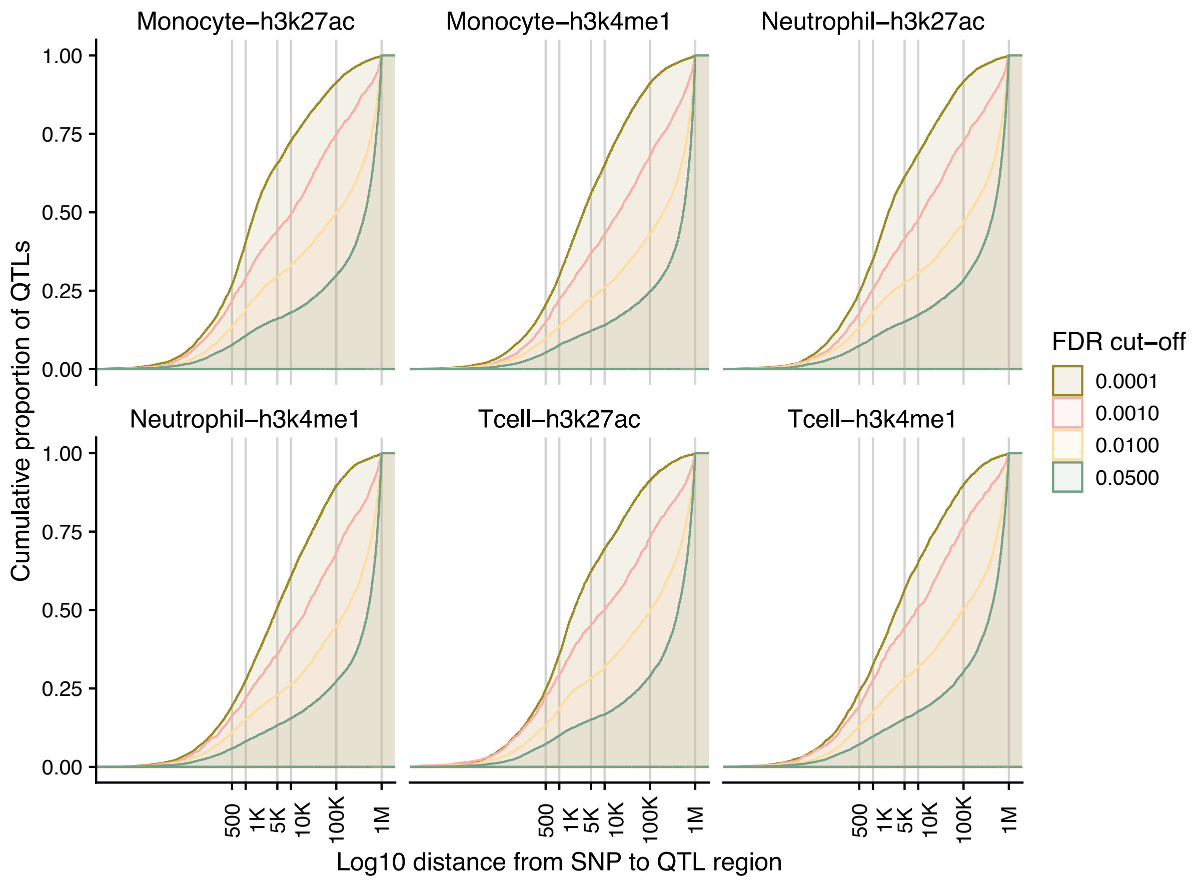


**Fig. S2 Histone mark QTL data shows that SNPs regulate peaks that are significant distances away**. The distance from the SNP to the associated histone mark peak for the six histone mark quantative trait loci (hQTL) studies from BLUEPRINT Phase 2 are shown at increasing FDR cut-offs. Here, to avoid double counting due to linkage disequilibrium (LD), SNPs were aggregated to those with the lowest p-value for each LD block based on Pickrell LD blocks. The y-axis is the cumulative proportion of interactions and the x-axis is the $\log_{10}$ distance with a grey vertical lines increasing numbers of base-pairs. At all FDR cut-offs, for all hQTL studies, we observe high proportions of interactions (between 20%-75%) at distances above 10,000 base-pairs. This goes against a null hypothesis that SNPs only regulate the epigenome by local motif disruption and highlights the need for models which can account for these distal interactions when predicting epigenetic profiles.

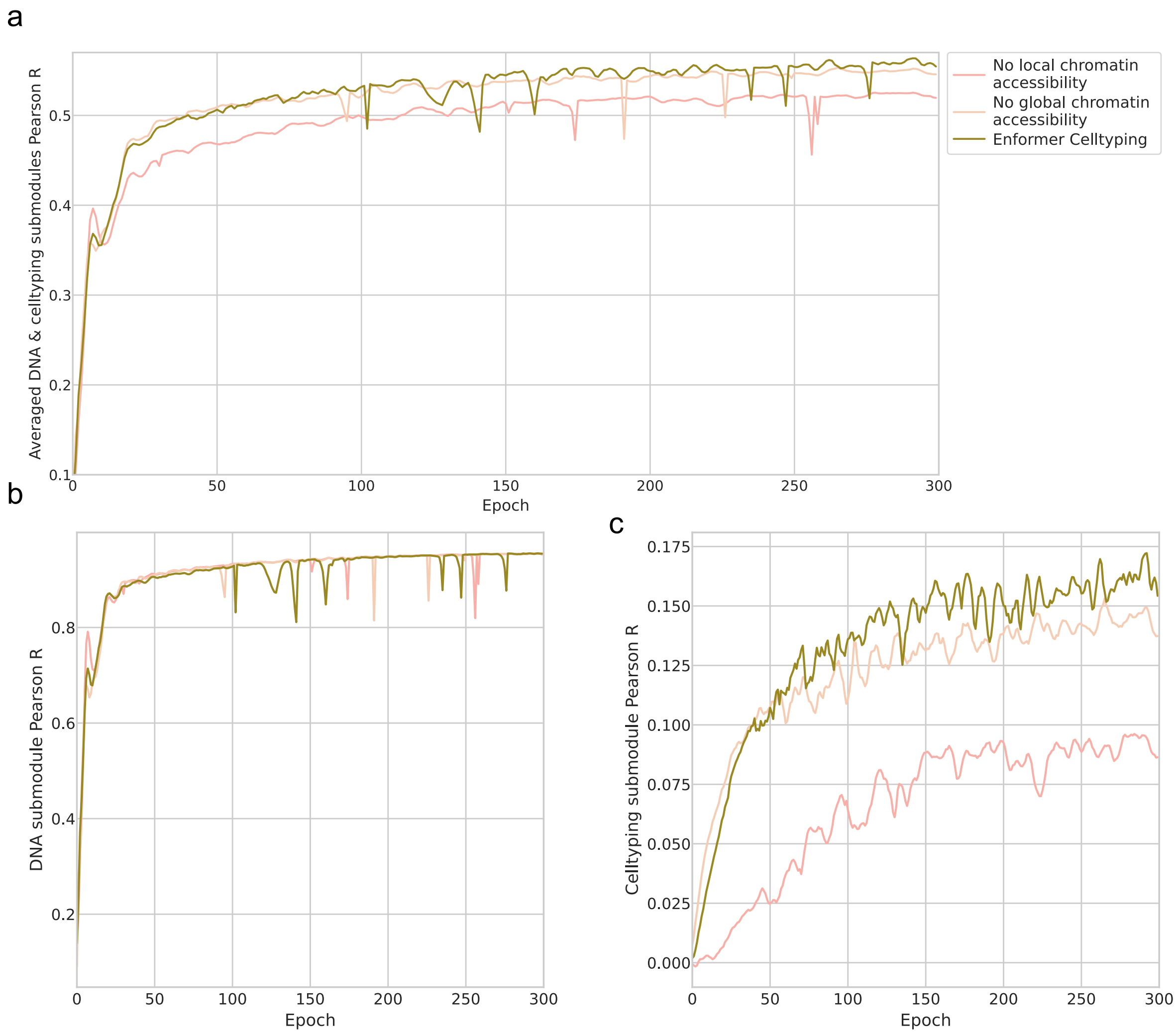


**Fig. S3 Using global and local chromatin accessibility information improves the prediction of cell type-specific histone mark signals**. We measured the performance of Enformer Celltyping on the validation set for the pre-training stage over 300 training steps (batch size 128) for both global and local chromatin accessibility information (green), no local chromatin accessibility (pink) and no global chromatin accessibility (beige). (a) Averaged Pearson R performance for both the DNA and celltyping submodules, (b) DNA submodule – note this is a negative control, these should be the same since the chromatin accessibility only effects the celltyping submodule and (c) the celltyping submodule – where this should have the largest effect. All Pearson R values are performance on the validation dataset.

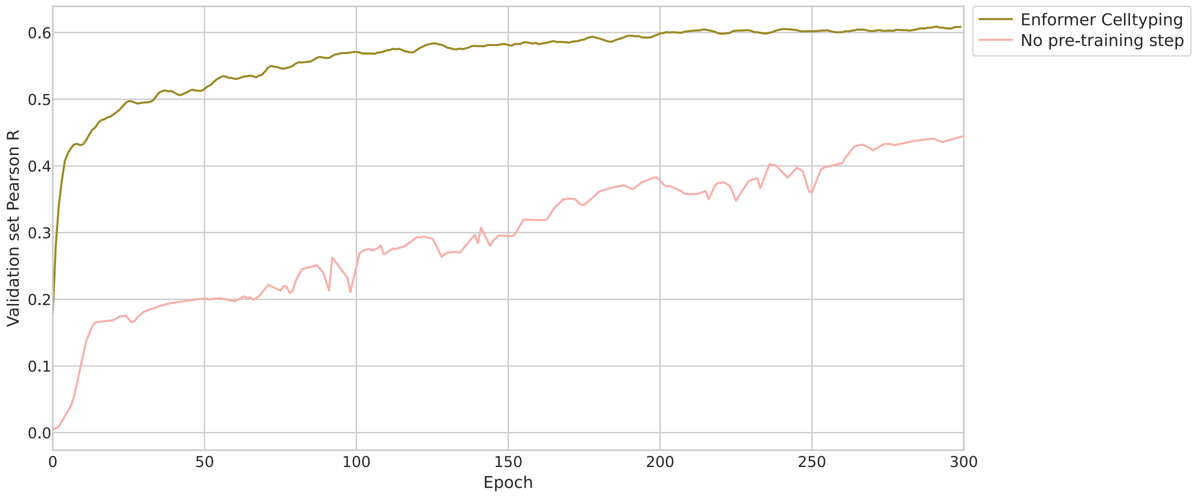


**Fig. S4 Pre-training drastically improves Enformer Celltyping's performance**. We measured the performance of Enformer Celltyping on the validation set for the full training stage over 300 training steps (batch size 128) comparing the full Enformer Celltyping model (green) against a version of Enformer Celltyping for which a pre-training stage was not performed (pink). Pearson R values are performance on the validation dataset.

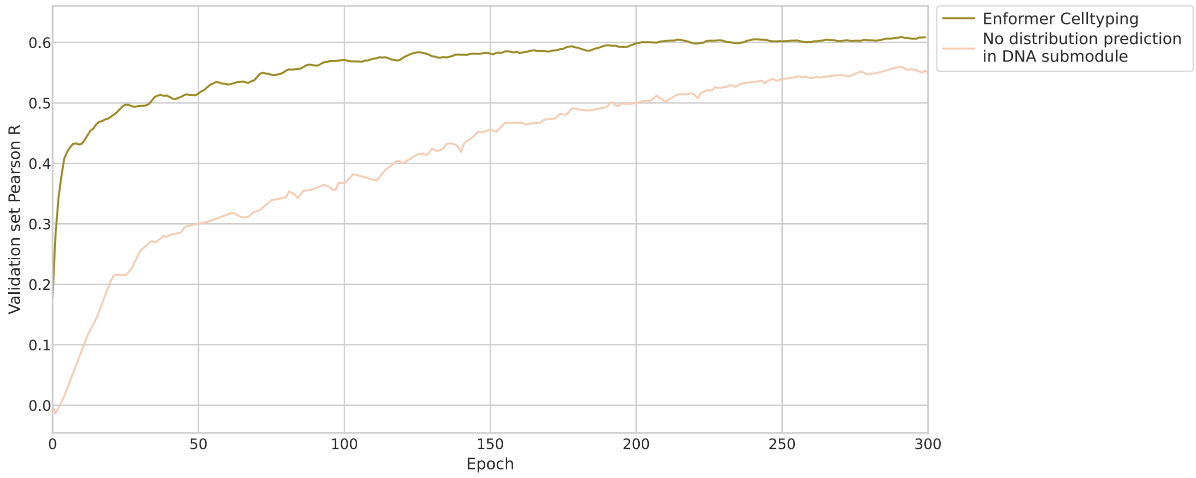


**Fig. S5 Predicting the distribution of signals as well as the average with the DNA module during pre-training improves Enformer Celltyping's performance**. We measured the performance of Enformer Celltyping on the validation set for the full training stage over 300 training steps (batch size 128) comparing the full Enformer Celltyping model (green) against a version of Enformer Celltyping for which the DNA module only predicted the average expression, as opposed to the average and distribution, in the pre-training stage (beige). Pearson R values are performance on the validation dataset.

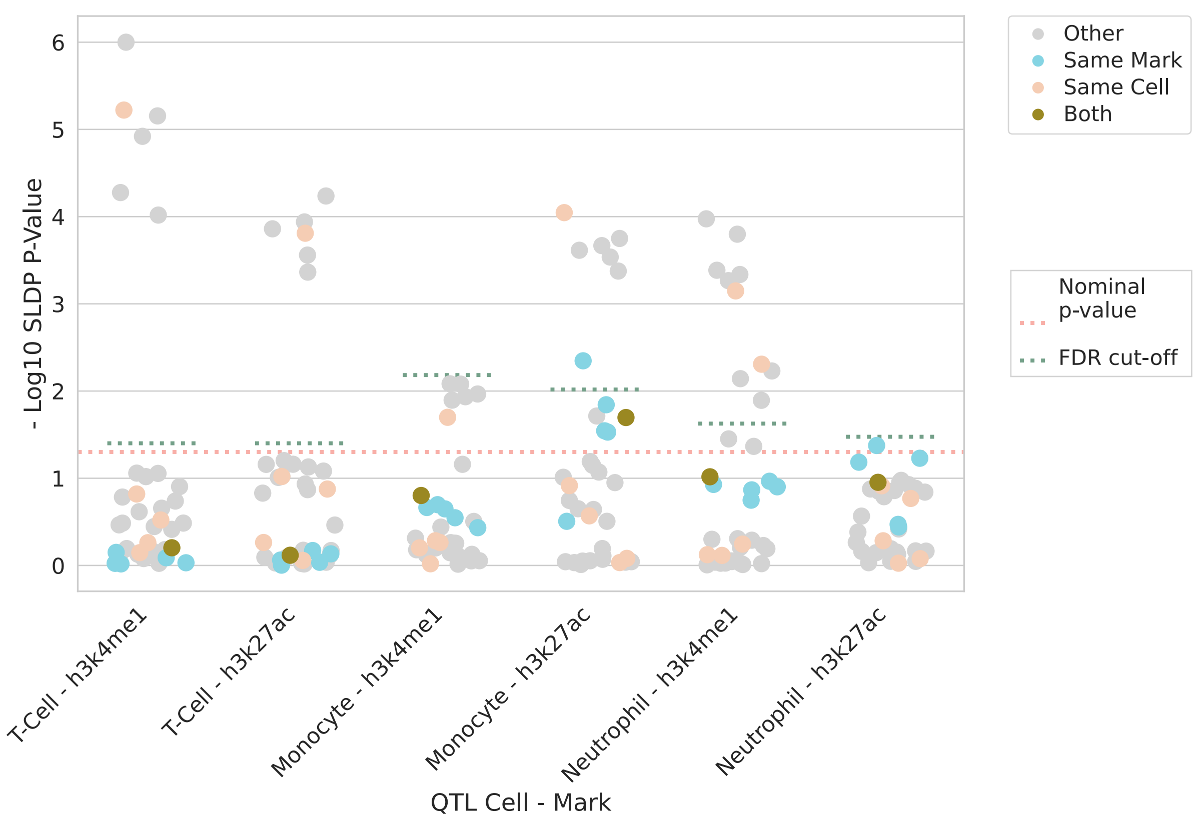


**Fig. S6 Enformer Celltyping's genetic effect predictions are note noticeably improved by filtered out QTLs associated with downregulating peaks which are missed in predictions for the major allele.** Statistical significance (y-axis) of SLDP genome-wide concordance between Enformer Celltyping's genetic variant predictions and measured hQTL effect sizes. Here, any SNP to histone mark binding hQTL entries were removed if it caused a decrease in binding and Enformer Celltyping did not predict the binding on the major allele (on the DNA sequence without the genetic variant). The x-axis depicts the six Blueprint phase 2 hQTL studies and the cell-histone mark predictions are coloured by their relationship to the hQTL study: 'Both' – both the histone mark and the cell predicted in were the same as the hQTL study, 'Same Mark' – the mark matched, 'Same Cell' – the cell matched and 'Other' – neither the cell nor mark matched. If there was perfect agreement between our model and the hQTL studies, all 'Both' entries would be significant. A nominal p-value (pink dashed line) and the cut-off after Benjamini and Hochberg correction based on the minimum, non-significant p-value, corrected at the level of each hQTL (Turquoise dashed line) are both added.

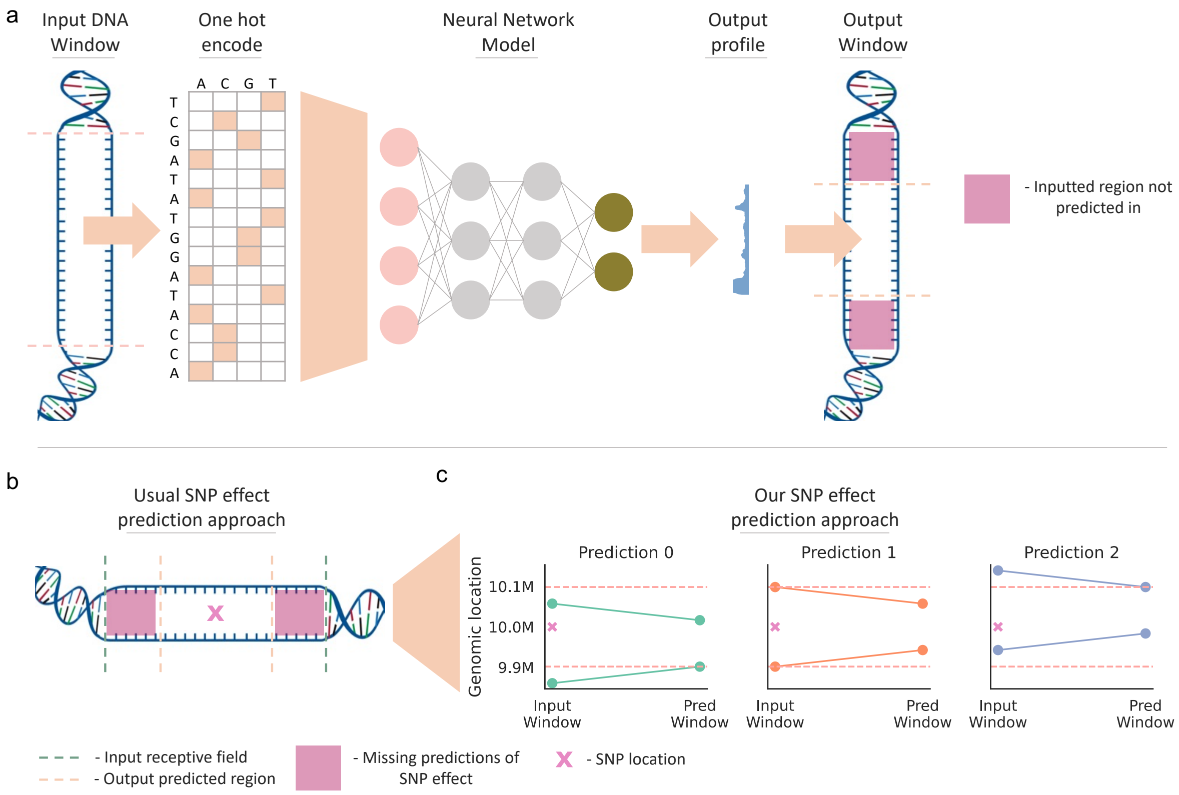


**Fig. S7 Schematic showing our strategy to capture distal effect of SNPs, by predicting each SNP multiple times, sliding the input window around the SNP location**. Genomic deep learning models tend to predict profiles for only the centre proportion of the genomic location relating to the DNA input to avoid predicting on edge regions which have less neighbouring genomic information. This funnel effect is shown in (a). Thereafter, common practice when predicting the effect of SNPs (i.e. *in silico* mutagenesis) with these models, is to centre the SNP on the input window (b). However, this approach misses distal predictions of SNP effects, not using the full receptive field of the model. We propose an alternative strategy to capture the effect of the SNP across the full receptive field by predicting each SNP multiple times, sliding the input window around the SNP location, e.g. move the relative SNP location towards the edges, or closer to the centre (c). For Enformer Celltyping, this requires three predictions (c): A prediction made with the SNP in the centre of the input window (‘Prediction 1’) to capture the nearby effect of the SNP and sliding the input window to make two further predictions (‘Prediction 0’ and ‘Prediction 2’) to capture distal SNP predictions.

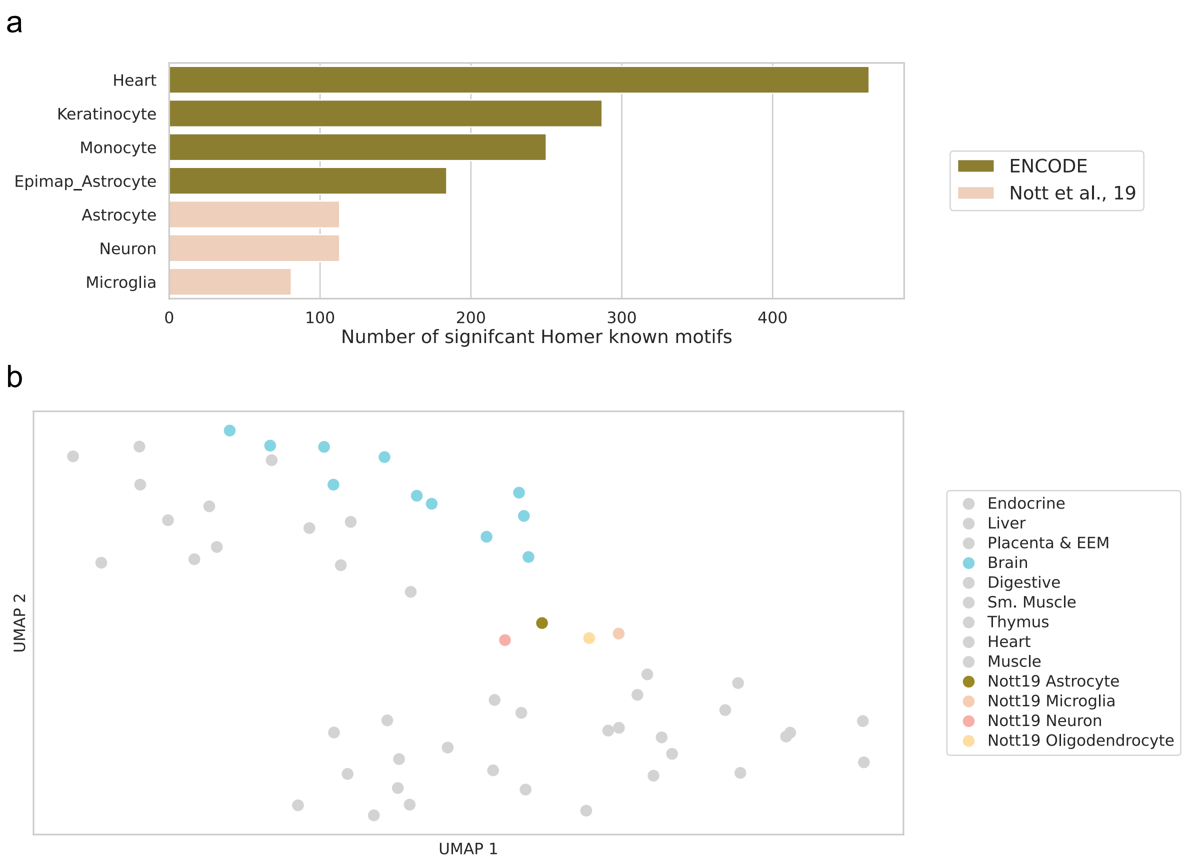


**Fig. S8 Enformer Celltyping's embedded global chromatin accessibility signal appears biased towards ENCODE samples.** (a) Gives the number of significant known motifs identified with Homer for each of the six cell types tested. The genomic regions for the motif analysis were derived from the histone mark peaks that had the highest contribution from the global chromatin accessibility signal for each cell type, identified by dropping the global signal inputted to the model and measuring the change in predicted binding. The cell types are coloured by their source. (b) A UMAP projection of Enformer Celltyping's global chromatin accessibility embeddings. Both training cell types and the four isolated cortical brain cell types from Nott et al. are included (the later are coloured). The other training cell types related to the brain (all bulk tissue, not isolated cells) are also coloured.

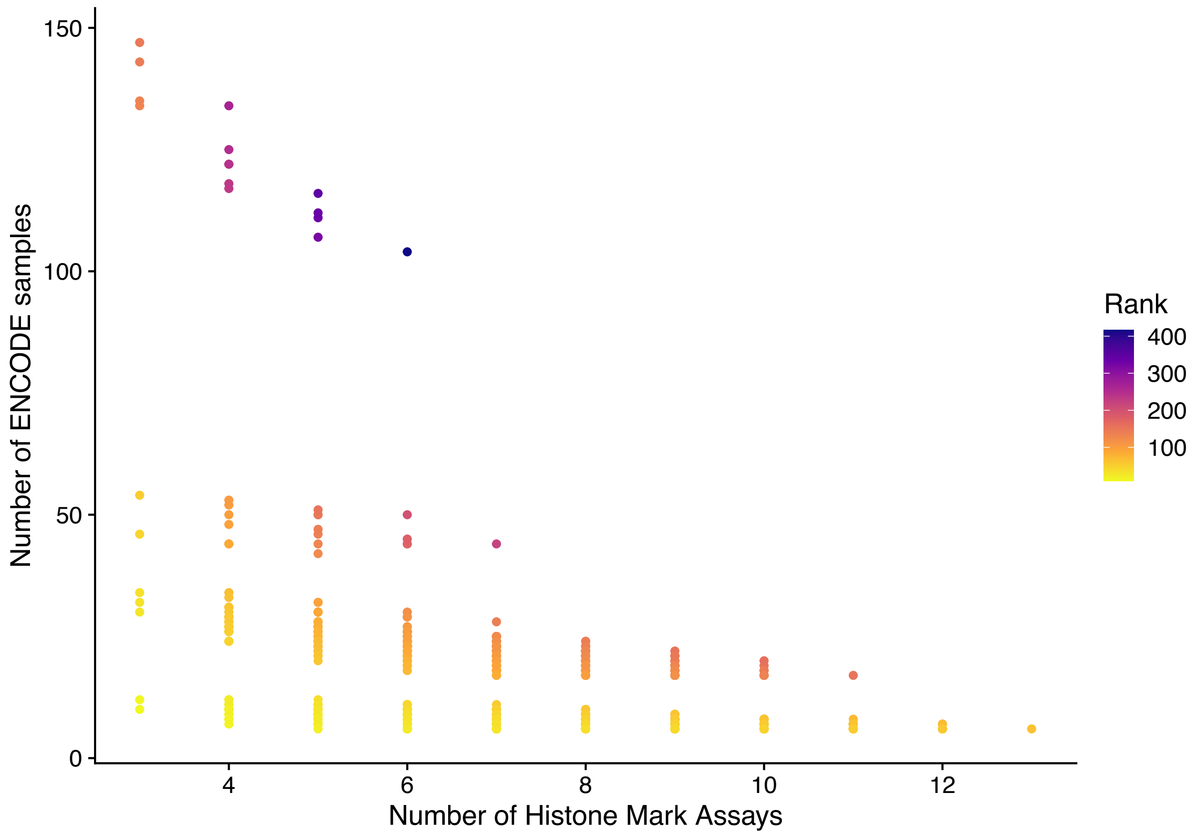


**Fig. S9 EpiMap trade-off between the number of samples and the number of assays**. The histone mark data available from EpiMap is shown with the number of samples (cell/tissue type) along the y-axis at each of the different number of histone mark assays along the x-axis. When choosing the number of assays to include in the model's predictions, the aim was to maximise both the number of assays but also the number of samples. The colour, which is the product of these numbers, of each position gives a rank which takes both into account. Note that the number of samples exclude our 3 validation cells (CD14+ monocytes, CD16+ neutrophils and naive CD4+T cells) and any related cell types.

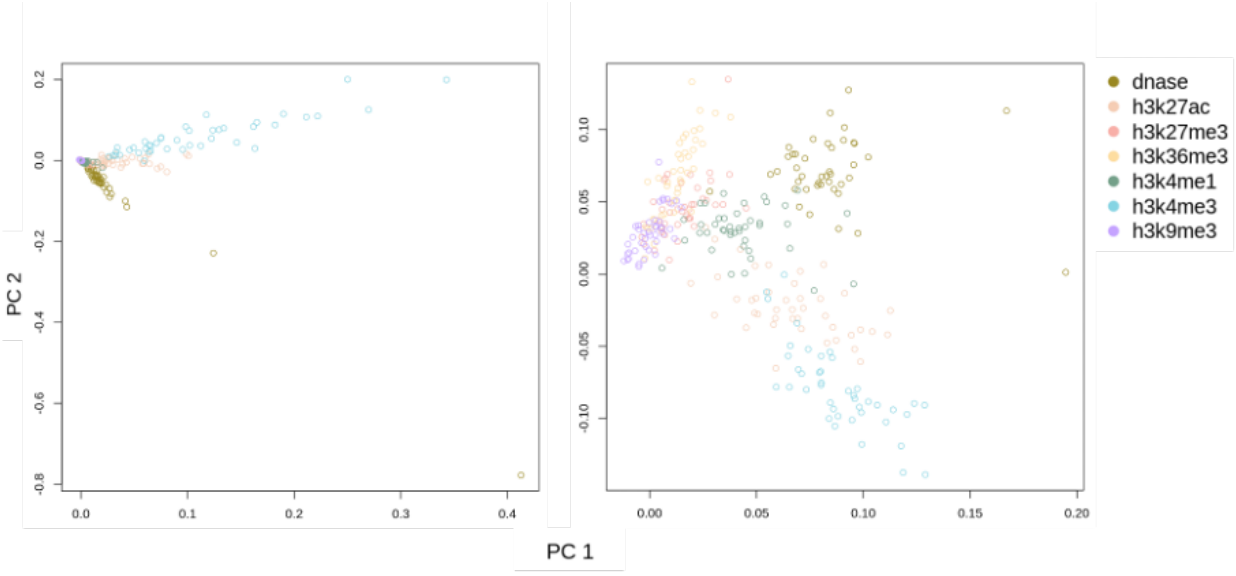


**Fig. S10 Arcsinh-transformed epigenetic signals for same marks show groupings**. Principal component analysis (PCA) plot of the first two principal components without (left) and with (right) arcsinh-transform based on the EpiMap training samples. Note that the data is not scaled here (a usual step in PCA) to replicate the data a model would be trained on. The arcsinh-transformed data shows clear separation between assay types, something which is not present with the raw data, highlighting arcsinh-transform's benefit. Note that DNase-Seq assayed were included here to show chromatin accessibility differentiation although they were not used in the model training approach.
